# Comparative Analysis of Primary and Liver Fibroblasts Reveals MET as a Potent Target in Pancreatic Cancer Metastasis

**DOI:** 10.1101/2025.09.06.674373

**Authors:** Rima Singh, Natalie Yousefian, Walker M. Allen, Cecily Anaraki, Alica K. Beutel, Sabrina Calderon, Ian M. Loveless, Oliver G. McDonald, Jennifer P. Morton, David Imagawa, Zeljka Jutric, Thomas F. Martinez, Nina G. Steele, Christopher J. Halbrook

**Affiliations:** Department of Molecular Biology and Biochemistry, University of California Irvine, Irvine, CA 92697, USA; Department of Internal Medicine I, Ulm University Hospital, Ulm, Germany; Pancreatic Cancer Center, Henry Ford Health System, Detroit, MI 48202, USA; Department: Pathology and Laboratory Medicine, University of Miami, Miami, FL; Cancer Research UK Scotland Institute, Glasgow G61 1BD, UK; Institute of Cancer Sciences, University of Glasgow, Glasgow G61 1BD, UK; Department of Surgery, University of California Irvine, Orange, CA 92868, USA; Chao Family Comprehensive Cancer Center, University of California Irvine, Orange, CA 92868, USA; Department of Pharmaceutical Sciences, University of California Irvine, Irvine, CA 92697, USA; Department of Biological Chemistry, University of California Irvine, Irvine, CA 92697, USA; Division of Digestive Diseases, Department of Internal Medicine, University of Cincinnati, Cincinnati OH 45212, USA; Institute for Immunology, University of California Irvine, Irvine, CA 92697, USA

## Abstract

The tumor microenvironment drives many malignant features of pancreatic ductal adenocarcinoma (PDAC). The fibroblasts within pancreatic tumors promote tissue remodeling, immune suppression, and resistance to therapy. However, the interactions between stromal populations and pancreatic cancer cells are less understood in the liver, the most frequent site of PDAC metastasis. To address this, we employ single cell transcriptomics to compare primary pancreatic vs. liver PDAC lesions. Here, we identify the expression of hepatocyte growth factor (*HGF*) in fibroblasts and its receptor *MET* in cancer cells are both markedly increased in the PDAC liver niche. Using functional assays, we validate that mitogenic MET signaling is activated in PDAC cells by liver-derived fibroblasts. Importantly, the inhibition of MET signaling leads to reduced tumor growth in immune competent mouse models. Collectively, our data demonstrates that liver stromal-epithelial crosstalk networks engage in signaling pathways distinct from primary pancreatic tumors, highlighting opportunities to develop new treatments for metastatic disease.

## Introduction

Pancreatic Ductal Adenocarcinoma (PDAC) remains one of the deadliest major cancers(1). The diversity of cell populations within pancreatic tumors have been shown to provide numerous avenues supporting cancer growth and survival (2). Cancer associated fibroblasts (CAFs) often constitute the majority of overall cellularity within the dense stroma, driving an intense fibroinflammatory reaction (3). This characteristic tissue remodeling by CAFs impairs drug and nutrient diffusion within primary PDAC tumors (4,5), but also acts to restrain metastasis (6–8). Accordingly, there have been extensive efforts to understand the varied, occasionally paradoxical, roles of CAFs in the pancreatic tumor microenvironment.

Early efforts to understand the stroma during pancreatic tumorigenesis predominantly identified CAFs as an expansion of a pancreatic stellate cell population based on features reminiscent of liver stellate cells, such as vitamin A droplet accumulation (3). However, a compendium of work has uncovered a remarkable heterogeneity among CAFs within pancreatic tumors (2). Recent lineage tracing of *Fabp4*^+^ cells suggests that only a small, but functionally important, subset of pancreatic CAFs are derived from a stellate cell origin (9). In contrast, up to half of the PDAC CAF population can arise from *Gli1^+^*pancreas resident fibroblasts(10). Regardless of origin, pancreatic CAFs have been shown to have functional phenotypes that include myofibroblast CAFs (myCAFs) responsible for stromal remodeling (11), inflammatory CAFs (iCAFs) that modulate immune suppression (11), antigen-presenting CAFs (apCAFs) characterized by high MHC II expression (12,13), and senescent CAFs that have been shown to have pro-tumorigenic role (14). Pancreatic CAFs can also be separated into functionally distinct roles by CD105 expression (15). Moreover, CAF heterogeneity fuels spatially distinct immune hot “reactive” and matrix rich “deserted” sub-tumor microenvironments that often co-exist within individual pancreatic tumors (16).

Importantly, the volume and diversity of CAFs allow for a multitude of avenues to dynamically interact with PDAC cells in a way that actively supports tumor growth. For example; IGF1, GAS6, and LIF released from CAFs provide paracrine signals to PDAC cells that support numerous pro-tumor pathways (17,18). Further, CAFs can directly supply metabolites including lipoproteins (19), amino acids (20), nucleotides (21), and glycosylation intermediates to support PDAC metabolism which is notoriously dysregulated by poor nutrient availability within the pancreatic tumor microenvironment (22–24). Similarly, Netrin G1^+^ CAFs support PDAC progression through the modulation of glutamine metabolism while suppressing an anti-tumor immune response (25).

Many of these crosstalk pathways between pancreatic CAFs and cancer cells have been investigated as therapeutic targets to improve the poor prognosis of PDAC patients. However, only 20% of patients diagnosed with this deadly malignancy present with early stage disease where surgical resection, combined with (neo)adjuvant chemotherapy, is the preferred treatment approach (26). In contrast, 80% of PDAC patients present with metastatic disease, with the liver being the most frequently colonized secondary organ (27). In addition, the liver also represents the most frequent site of distant recurrence in patients who have undergone primary tumor resection. The liver harbors its own diverse resident fibroblast populations (28). While some of these liver-resident fibroblasts exhibit similarity to some that are found in the pancreas, such as hepatic stellate cells, CAFs that arise in the PDAC liver niche interact with different cell populations encountered in their microenvironment as compared to CAFs found in primary pancreatic tumors (29,30). Consequently, approaches developed to target stromal-cancer crosstalk that have been derived from studying pancreatic CAFs may or may not retain therapeutic benefit in metastatic disease.

To address this, we set out to contrast the phenotype of CAFs in liver tumors vs. pancreatic tumors derived from PDAC. Leveraging single cell RNA sequencing of murine tumors derived from the same cancer cells, we identify that the majority of the CAFs isolated from liver PDAC do not correspond to transcriptomic profiles that describe CAFs present in primary PDAC. Using the receptor-ligand pairing tool CellChat, we identify a putative *HGF*-*MET* crosstalk axis that is exclusive to liver CAFs and PDAC cells. Isolating primary human fibroblasts, we functionally demonstrate that liver-derived fibroblasts release HGF that can activate mitogenic signaling pathways in PDAC cells. Finally, we demonstrate that both pharmacological and genetic MET targeting markedly impair the growth of liver PDAC in immune competent mice. Collectively, these data demonstrate that programming of cancer-associated fibroblasts is influenced heavily by the tissue from which they arise. This leads to potential opportunities to develop new approaches tailored to target stromal-cancer crosstalk in an organ-specific fashion.

## Results

### Fibroblast populations in the liver are expanded in the PDAC metastatic tumor niche

To compare the changes in the stromal compartment between the pancreas and liver in response to the presence of PDAC, we performed histology on matched tissues from the autochthonous Kras^+/LSL-G12D^;Trp53^+/LSL-R172H^;Pdx1-Cre (KPC) murine PDAC model vs. wildtype mice. In both the liver and the pancreas, PDAC lesions are accompanied by an expansion of the stroma within tumor region (**Fig. 1a**). Further, by immunostaining for the pan-fibroblast marker Platelet-derived Growth Factor Receptor (PDGFR) we observe that fibroblasts are readily present across the normal liver and that their numbers are markedly expanded in PDAC metastatic lesions, similar to the stromal expansion seen in primary pancreatic tumors vs. normal pancreas (**Fig. 1b,c**). In addition, using in situ hybridization for the human fibroblast marker Decorin (*DCN)* in human PDAC liver biopsy samples, we readily observe fibroblasts across nearby normal liver tissues that remain closely associated with the epithelial cells (**Fig. 1d**). Collectively, these data suggest that the abundant fibroblast populations in the liver have potential to serve many of the pro-tumorigenic roles that have been previously ascribed to pancreatic CAF populations.

**Figure 1:**
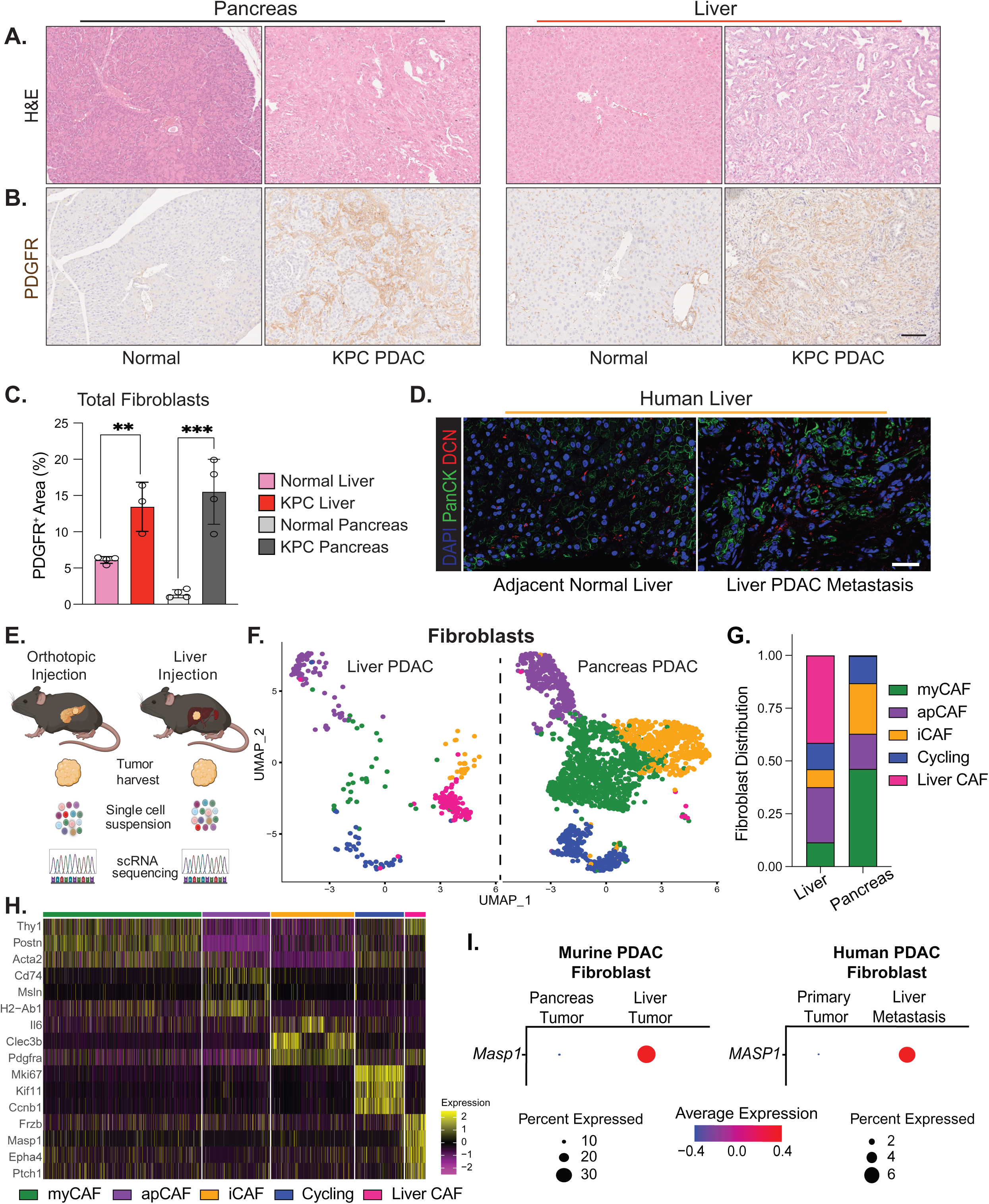
Stromal expansion occurs in PDAC liver lesions with distinct fibroblasts from pancreatic tumors. **A.** Hematoxylin and Eosin (H&E) staining of normal tissue pancreas or liver tissues from wildtype mice compared with PDAC primary and liver metastatic tumors from KPC mice. **B.** Representative immunohistochemistry of the pan-fibroblast marker PDGFRβ in tumor tissue vs. normal liver and pancreas. **C.** Quantification of PDGFRβ^+^ area in (**B**) (n=3 liver samples, n=4 pancreas). **D.** Fluorescent ISH staining for the fibroblast marker Decorin (DCN, red) coupled with IF for the epithelial marker Pan-Cytokeratin (PanCK, green) in primary human PDAC and liver metastasis. **E.** Schematic illustration of experimental design for single cell RNA sequencing comparing pancreas vs. liver PDAC tumors. Here, syngeneic KPC cells are injected in mice orthotopically to form tumors in the pancreas or via hemi-spleen injection to seed tumors in the liver and processed for sequencing. **F.** Uniform manifold approximation and projection (UMAP) plot showing the fibroblasts populations present in either pancreas or liver PDAC lesions. **G.** Ratio of the distribution of cells between myCAF, apCAF, iCAF, cycling, or Liver CAF sub-populations in either pancreas or liver PDAC tumors. **H.** Top 5 heatmap of genes that separate fibroblast sub-populations. **I**. Dot plot representation of the gene expression of the Liver CAF marker *MASP1* between fibroblast populations present in pancreas or liver lesions in murine or human PDAC. Scale bar = 100µM. Error bars are mean ±SD, ** P ≤ 0.01; *** P ≤ 0.001 by two-tail student’s t test.

### Single-cell transcriptomics show liver CAFs are distinct from pancreatic CAF populations

Previous studies have consistently found that single cell sequencing of pancreatic biopsies underrepresent fibroblast populations in samples obtained from tumor resections (31). As liver PDAC metastasis specimens are nearly always obtained via biopsy, even large collections of data such as our Human Pancreatic Cancer Single-Cell Atlas have complicated studying liver PDAC CAFs in detail (32). Finally, the genetic diversity across patients with very few paired primary and liver metastatic lesions limits the ability to compare stromal cells between tumor sites.

Accordingly, we leveraged comparative murine allograft models by either injecting syngeneic KPC FC1245 PDAC cells into the liver of C57BL/6J mice via a hemi-splenic injection method or injected the same PDAC cells orthotopically into the pancreas and harvested the tumors on day 18 and day 15, respectively (**Fig. 1e**). To gain insight into global programming of the pancreas vs. liver PDAC lesions we performed 10X chromium single cell RNA sequencing on the total cell population dissociated from microdissected tumors. Using the Louvain algorithm, we clustered the cells from both tumor sites together in an unbiased manner (**Supplemental Fig. 1a**). Major cell types were annotated using canonical markers (**Supp. Fig. 1b,c**) (33) allowing us to identify eleven different populations of epithelial, immune, and stromal cells in these tumors. These populations were largely represented across both tumor sites (**Supplemental Fig. 1d**), however the fractional distribution of the populations captured varied (**Supplemental Fig. 1e**). Thus, while single cell transcriptomics are not necessarily a quantitative measure of absolute cell numbers, these trends can nonetheless inform some of the potential differences in the immune infiltration in the pancreatic vs. liver PDAC tumor microenvironment.

Importantly, we captured a sufficient population of fibroblasts between the pancreatic and liver PDAC lesions to enable further subclustering. Within our fibroblast object, we identified 11 clusters (**Supplemental Fig. 2a**) that largely mapped to the previously described myCAF, iCAF, and apCAF transcriptional subtypes that have previously been described. However, beyond a cluster of clearly proliferating fibroblasts, one population was clearly distinct and did not share transcriptional overlap with other reported pancreatic CAF subtypes such as senCAFs. Splitting the object by tumor site we find that this population is exclusively present in PDAC liver tumors (**Fig. 1f**), and these liver CAFs account for approximately half of the non-proliferating stromal object (**Fig. 1g**). Using transcriptional characterization of CAF sub-populations by top gene expression, we found previously established markers of CAF populations such as *Il6* in iCAFs, *Acta2* in myCAFs, and *Msln* in apCAFs, and additionally, we were able to define putative marker genes for liver CAFs that include *Masp1, Frzb, Epha4, and Ptch1* (**Fig. 1h**). Of these, we observed that the specificity of *Masp1* expression in murine liver CAFs vs. pancreatic CAFs is mirrored in the CAFs found in human liver metastasis vs. primary CAF populations in our compendium of human single cell data (**Fig. 1i**). Finally, we validated MASP1 expression is indeed specific to liver vs. pancreatic CAFs in murine and human PDAC using immunostaining (**Supplemental Fig. 2b-d**).

Collectively, these data establish that the stromal populations associated with liver PDAC lesions are distinct from those that have been extensively characterized in primary pancreatic tumors. Accordingly, we inferred that liver CAFs will also employ mechanisms to shape the microenvironment of the metastatic niche unique from those that have been identified by studying pancreatic CAFs.

### PDAC cells display preferential metabolic programming in liver vs pancreas tumors

To determine how liver CAFs communicate with cancer cells, we proceeded to define the PDAC cell populations present between our syngeneic murine liver and pancreas tumors. Out of the initial 20 clusters, (**Supplemental Fig. 3a**) we found 4 main populations that displayed clear themes by pathway analysis of upregulated genes in each cluster (**Fig. 2a, Supplemental Table 1**) and two minor populations with unclear functions. Further investigating top genes associated with these pathways, we found that the two largest populations separated based on different metabolic preferences: glycolysis, or redox metabolism centered around protection from oxidative stress that we termed “Redox” (**Fig. 2b-d, Supplemental Fig. 3b**). The other two main populations were either centered on proliferation, or an enrichment of genes associated with extracellular interaction that we termed “ECM-related”. Given their unclear function, we termed the remaining minor populations “Epithelial 5”, and “Epithelial 6”.

**Figure 2:**
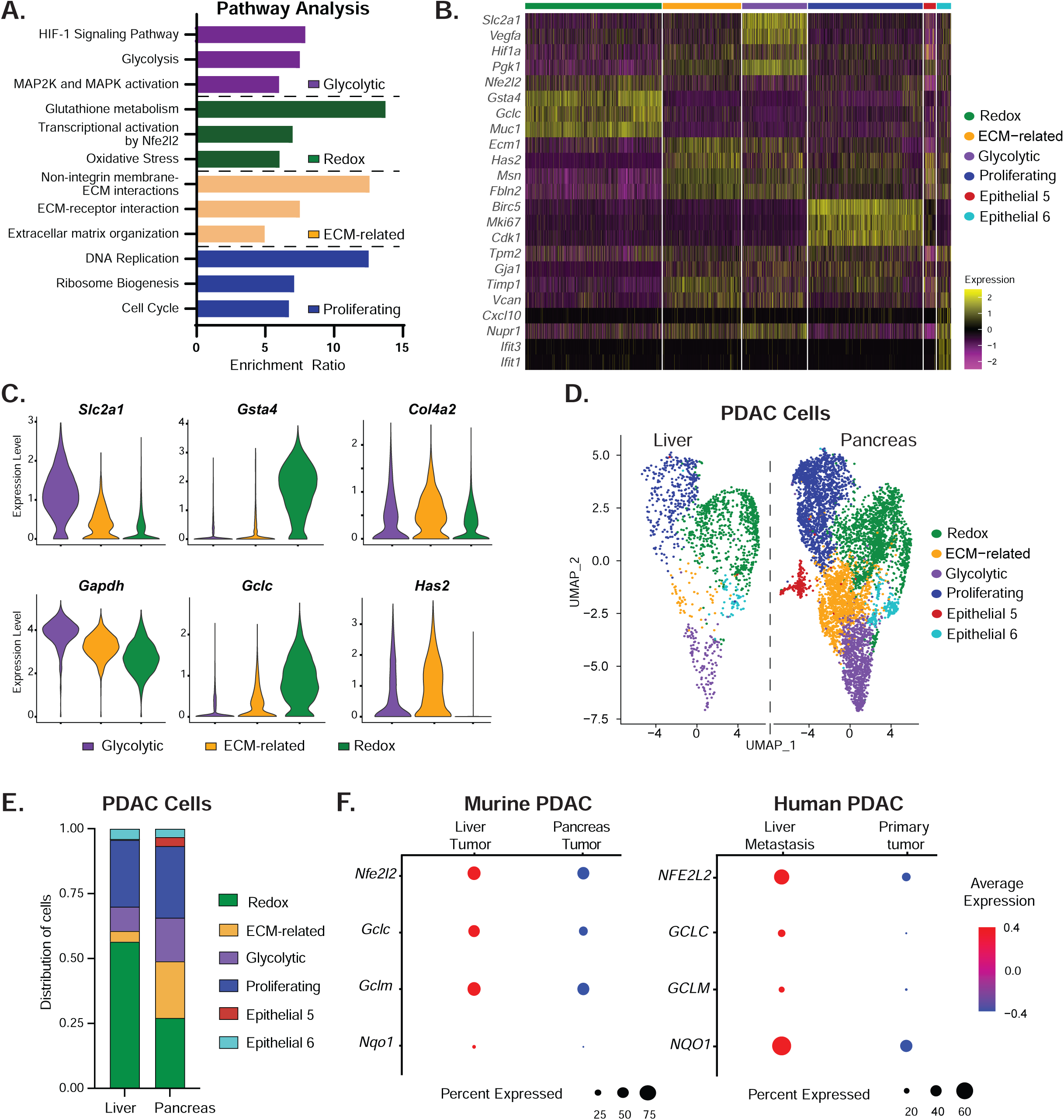
PDAC cells display metabolic heterogeneity and differential preferences between pancreas and liver tumors. **A.** Analysis of overrepresented genes in the main PDAC clusters grouped in distinct biological pathways centered on glycolysis, redox metabolism, extracellular matrix-related genes, or proliferation. **B.** Heatmap of highly expressed genes within each PDAC subpopulation. **C.** Violin plots showing expression of representative genes that distinguish the main PDAC clusters including *Slc2a1* and *Gapdh* (Glycolysis), *Gsta4 and Gclc* (Redox), or *Col4a2* and *Has2* (ECM-related). **D.** UMAP and bar graph (**E**) representation of PDAC population distribution between pancreas and liver lesions. **F.** Dot plot representation of redox metabolism genes *NFE2L2, GCLC, GCLM*, and *NQO1* between liver and pancreas PDAC cells from human or mouse tumors.

Interestingly, we observed that with the exception of Epithelial 5, all the other PDAC cell populations can be observed in both pancreatic and liver allografts (**Fig. 2d**). However, the distribution of the PDAC subclusters between tumor sites varied dramatically (**Fig. 2e**). Here, we observed that the pancreatic tumors demonstrated fairly even heterogeneity, with glycolytic, redox, and ECM-related populations in approximately equal proportion among the non-proliferating cells. In contrast, the vast majority of PDAC cells in liver tumors show a gene expression program centered on redox metabolism, potentially due to higher availability of oxygen and nutrient availability in the liver vs. the hypoxic and nutrient-challenged pancreatic tumors.

To validate the metabolic preferences of PDAC cells residing in the liver vs. the pancreas, we compared the expression of the master redox protein NRF2 (*NFE2l2* gene) and canonical downstream NRF2 targets in our murine and published human datasets. Here, we observed a clear upregulation of *NFE2L2, GCLC, GCLM, and NQO1* in the liver PDAC populations compared to those isolated from pancreatic tumors (**Fig. 2f**). Interestingly, the preference of redox metabolism in liver resident PDAC cells is conserved regardless of basal, classical, or hybrid transcriptional subtype (**Supplemental Fig. 3c**) This data suggest that pancreatic cancer cells either reprogram their metabolism in response to the nutrient availability and organotrophic factors, or the initial establishment of tumors in these different environments selects for specific metabolic populations that that we have previously shown co-exist within PDAC (34,35).

### The HGF-MET signaling axis is specifically upregulated in the liver metastatic tumors

With defined stromal and cancer epithelial clusters, we next proceeded to probe the potential interactions between these populations within the liver PDAC lesions. To accomplish this, we leveraged the ligand-receptor interaction tool CellChat (36) to build a liver PDAC signaling network (**Fig. 3a**) and a pancreatic PDAC signaling network (**Supplemental Fig. 4a**). While there are key differences in the number of interactions and interaction strengths between different populations in liver vs. pancreatic tumors, stromal populations appear to be the dominate source of signal (**Fig. 3a**). Interestingly, in liver PDAC lesions, liver CAFs are the most active source of signaling to other populations (**Fig. 3b**). Focusing our analysis on unique signaling pathways showing significant communications and identified to be outputs from liver CAFs and input by PDAC populations, we identified that only the Hepatocyte Growth Factor (HGF) pathway fits these criteria (**Fig. 3c**). In contrast, several other pathways that derive signals from stromal populations are conserved between liver and pancreatic PDAC tumor networks including PTN and IL6 (**Supplemental Fig. 4b**).

**Figure 3:**
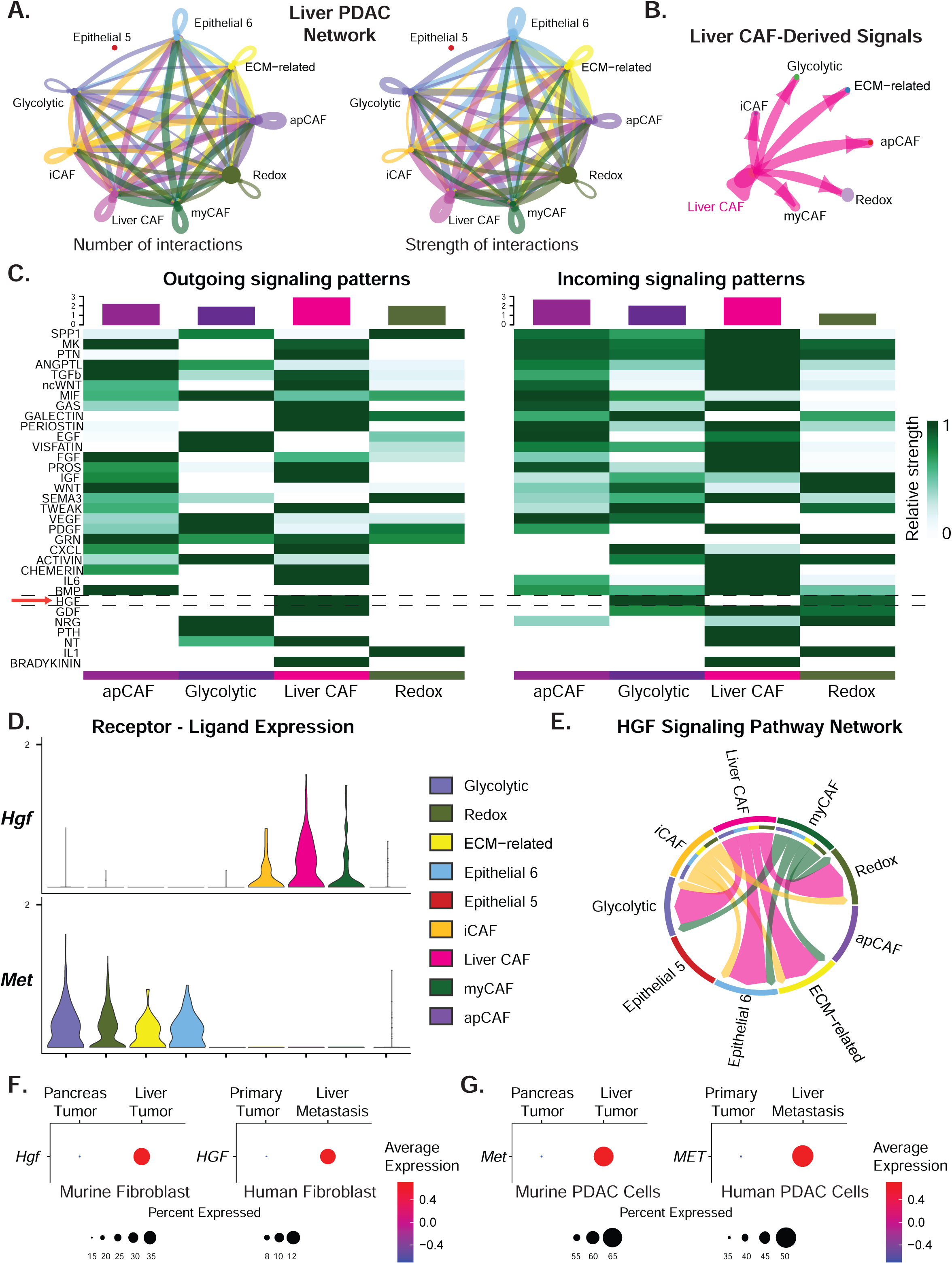
A paracrine HGF-MET signaling axis is upregulated in liver PDAC lesions. **A.** Circle plot depiction of the CellChat aggregated cell-cell communication network showing number of ligand-receptor interactions (left) and strength these interactions (right) between each indicated two populations in murine PDAC liver lesions. **B.** Isolated signaling from Liver CAFs to stromal and cancer populations in the liver niche. **C.** Top signaling pathways with the most outgoing or incoming signaling in PDAC liver, with HGF highlighted (red arrow). Violin plots showing gene expression of the ligand HGF (**D**) and the HGF receptor MET by stromal and PDAC sub-populations. **E.** The HGF signaling pathway visualized by chord diagram showing different CAF sub-populations signaling to the cancer cells, line width is proportional to interaction strength. **F.** Dot plot representation of the *HGF* gene expression between fibroblasts isolated from pancreas or liver PDAC lesions in mouse and human tissues. **G.** Dot plot representation of the *MET* gene expression between cancer cells isolated from pancreas or liver PDAC lesions in mouse and human tissues.

HGF, also known as Scatter Factor, is produced by mesenchymal cells, including both pancreatic and liver CAF populations (29,30,37). HGF is the only known ligand to the tyrosine kinase receptor MET. Using the Seurat wrapper function plotGeneExpression, we validated that liver CAFs are by far the most abundant source of *Hgf* expression among all the populations, and the expression of the *Met* receptor is exclusive to the PDAC populations (**Fig. 3d**). Further visualizing the distribution of the HGF pathway across these cell populations by chord diagram, we observe that liver CAFs are indeed the primary sender of the HGF signal to multiple epithelial clusters (**Fig. 3e**) whereas myCAFs and iCAFs only make minor contributions. Finally, we validated that the *HGF* ligand expression is upregulated in liver PDAC CAFs vs. primary pancreatic CAFs (**Fig. 3f**), and the expression of the *MET* receptor is upregulated in liver PDAC cells vs. pancreatic PDAC cells in both human and murine datasets.

### Human liver fibroblasts uniquely produce HGF to activate MET-ERK signaling in PDAC cells

To functionally examine the signaling promoted by liver vs. pancreatic fibroblast populations in human models, we established several primary fibroblast cultures from surgical resections that included a primary PDAC specimen (PDAC CAF), a liver-metastatic colorectal tumor (Liver CAF), and nearby normal liver tissue (Liver Fibroblast). We first collected RNA and performed bulk RNA-sequencing to compare these populations (**Fig. 4a**). Principal Component Analysis (PCA) showed distinct clustering of all three fibroblast populations (**Supplemental Fig. 5a**). Differential gene expression among the three groups showed more similarity between the two liver-derived fibroblasts than PDAC CAFs (**Supplemental Fig. 5b**). Unbiased clustering based on the z-scores of the top 50 differentially expressed genes which also included MASP1, further corroborated that fibroblasts tend to cluster on the organ of origin (**Supplemental Fig. 5c**).

**Figure 4:**
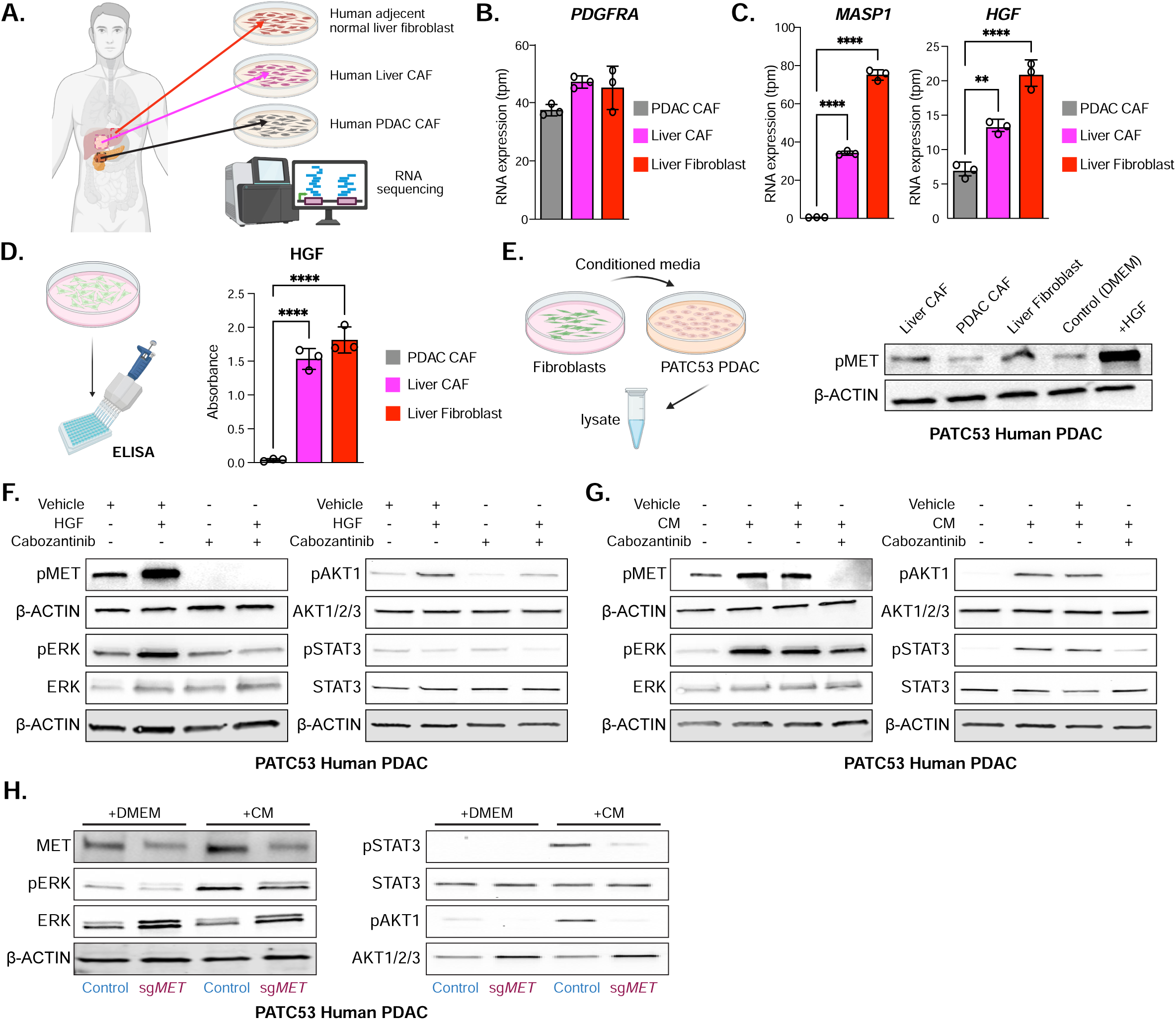
Human liver fibroblasts release HGF to activate mitogenic MET signaling in PDAC cells. **A.** RNA was isolated and sequenced from primary fibroblast cultures that were established from a resected liver colorectal metastasis (Liver CAF), the paired nearby normal liver tissue (Liver Fibroblast), or a primary PDAC tumor specimen (PDAC CAF). **B.** Quantification of the fibroblast marker gene *PDGFRA* across fibroblast populations. **C.** Quantification of Liver CAF marker *MASP1* and *HGF* gene expression across fibroblast populations. **D.** Quantification of HGF released by fibroblasts into conditioned media by ELISA. **E.** Western blot analysis of MET activation in human PDAC cell line PATC53 by a 3:1 ratio of conditioned media - fresh complete media from different fibroblast cultures as compared to HGF (1ng/mL) after 30 seconds. **F.** Western blot analysis of MET (30 seconds), STAT3, AKT, and ERK (15 minutes) in PATC53 PDAC cells treated with either HGF (1 ng/mL), the MET inhibitor cabozantinib (1µM) or combination. **G.** Western blot analysis of MET (30 seconds), STAT3, AKT and ERK (15 minutes) in PATC53 PDAC cells treated with either 3:1 liver fibroblast conditioned media:fresh media or fresh media alone or in combination with the MET inhibitor cabozantinib (1µM) or vehicle. **H.** Western blot analysis of MET, ERK, AKT, and STAT3 in sg*MET* vs. parental control PATC53 PDAC cells treated with a 3:1 ratio of liver fibroblast conditioned media:fresh media or fresh media. β-ACTIN was used as a loading control in all conditions. Error bars are mean ±SD; ** P ≤ 0.01; *** P ≤ 0.001; **** P ≤ 0.0001, by one-way Anova with Tukey post hoc.

Next, we compared specific gene expression between these fibroblast groups. Here, we found the expression of the fibroblast marker gene *PDGFRA* to be largely similar across both CAFs and the liver fibroblast cultures (**Fig. 4b**). In comparison, we observed that both the liver CAF marker MASP1 and expression of HGF are markedly upregulated in both liver-derived fibroblasts vs. PDAC CAFs (**Fig. 4c**). We next validated that the differences in *HGF* gene expression translated to a difference in protein production and release using an enzyme linked immunosorbent assay (ELISA) on conditioned media from the fibroblast cultures. As expected, we observe dramatically higher HGF production by both liver-derived fibroblast populations vs. PDAC CAFs (**Fig. 4d**).

To determine if the HGF released by liver-derived fibroblasts has a functional impact on human pancreatic cancer cells, we treated the PATC53 PDAC cell line with fibroblast conditioned media and examined MET activation (**Fig. 4e**). Indeed, we observed that the conditioned media from both human Liver-CAF and human adjacent normal liver fibroblasts resulted in increased MET phosphorylation vs. PDAC CAF conditioned media. Investigating potential downstream implications of MET activation in PDAC cells, we treated PATC53 cells with human recombinant HGF and observed an increase in ERK and AKT1 activation (**Fig. 4f**). This activation of MET, ERK, and AKT1 by HGF can be disrupted by treatment with the pharmacological MET inhibitor cabozantinib (XL-184). Importantly, we observe MET, ERK, AKT1 and STAT3 activation in PATC53 cells treated with liver fibroblast conditioned media, and these are also reduced by treatment with cabozantinib (**Fig. 4g**). Interestingly, the ERK and STAT3 activation by fibroblast conditioned media is only partially disrupted by MET inhibition, suggesting that other upstream pathways are likely being stimulated in parallel.

To further investigate the role of MET in activating mitogenic pathways, we genetically targeted MET using CRISPR editing. Here, we observe that partial loss of MET is sufficient to promote dramatic compensatory rewiring of mitogenic pathways downstream of MET in PATC53 cells (**Fig. 4h**). This includes a marked upregulation of the total protein levels of ERK, AKT, and to a lesser extent, STAT3 in sg*MET* PATC53 cells. Importantly, despite the increased overall protein levels, the activation of these pathways in response to treatment with liver fibroblast conditioned media is greatly diminished in sg*MET* vs. parental control. Again, we observe some ERK activation in sg*MET* relative to AKT, further suggesting the presence of other factors such as potential fibroblast-produced EGFR ligands may also be present(38).

Collectively, these data show that human liver-derived fibroblast populations are uniquely equipped to activate MET signaling in PDAC cells. MET activation drives multiple mitogenic signaling pathways that play centrally important roles in PDAC. Importantly, MET activation by stromal HGF can be targeted by clinically available pharmacological inhibitors such as cabozantinib.

### Genetic and Pharmacological MET inhibition impair growth of liver PDAC tumors in mice

Our data demonstrated that HGF production by liver stromal cells has functional consequences on pancreatic cancer cells in culture; this has the potential to be even more impactful in liver PDAC tumors given their increased MET expression we have observed on PDAC cells in the liver niche. Accordingly, we examined the impact of the MET inhibitor cabozantinib on liver PDAC tumors. To accomplish this, we established syngeneic KPC-FC1245 tumors in the livers of mice C57BL/6J through hemispleen injections or in the pancreas through orthotopic injections. After letting tumors establish for 6 days, we randomized mice onto treatment arms with either cabozantinib or vehicle via daily oral gavage (**Fig. 5a**). We monitored the mice every day and found no significant weight loss in the animals in this 16-day study (**Supplemental Fig. 6a**). On day 16, mice were sacrificed, and tissues were harvested. All livers harvested from control mice presented with clear macroscopic tumor lesions, whereas the majority of cabozantinib treated mice showed no obvious tumors in the liver (**Fig. 5b,c, Supplemental Figure 6b**). This was further validated by immunostaining for cytokeratin-19 (CK-19), a ductal epithelial marker that is a surrogate for PDAC cells. Here, we observed that the livers of vehicle treated mice were largely comprised of CK19^+^ tumor cells, whereas cabozantinib treated liver tissues showed either no tumors or a few small tumor masses within the liver tissue (**Fig. 5d,e**). In contrast, gross macroscopic tumors were present in the pancreas of all orthotopically injected mice regardless of treatment arm (**Fig. 5f,g, Supplemental Figure 6c**), and we observed that cabozantinib treatment modestly impairs primary final tumor mass vs. vehicle treatment (**Fig. 5h**).

**Figure 5:**
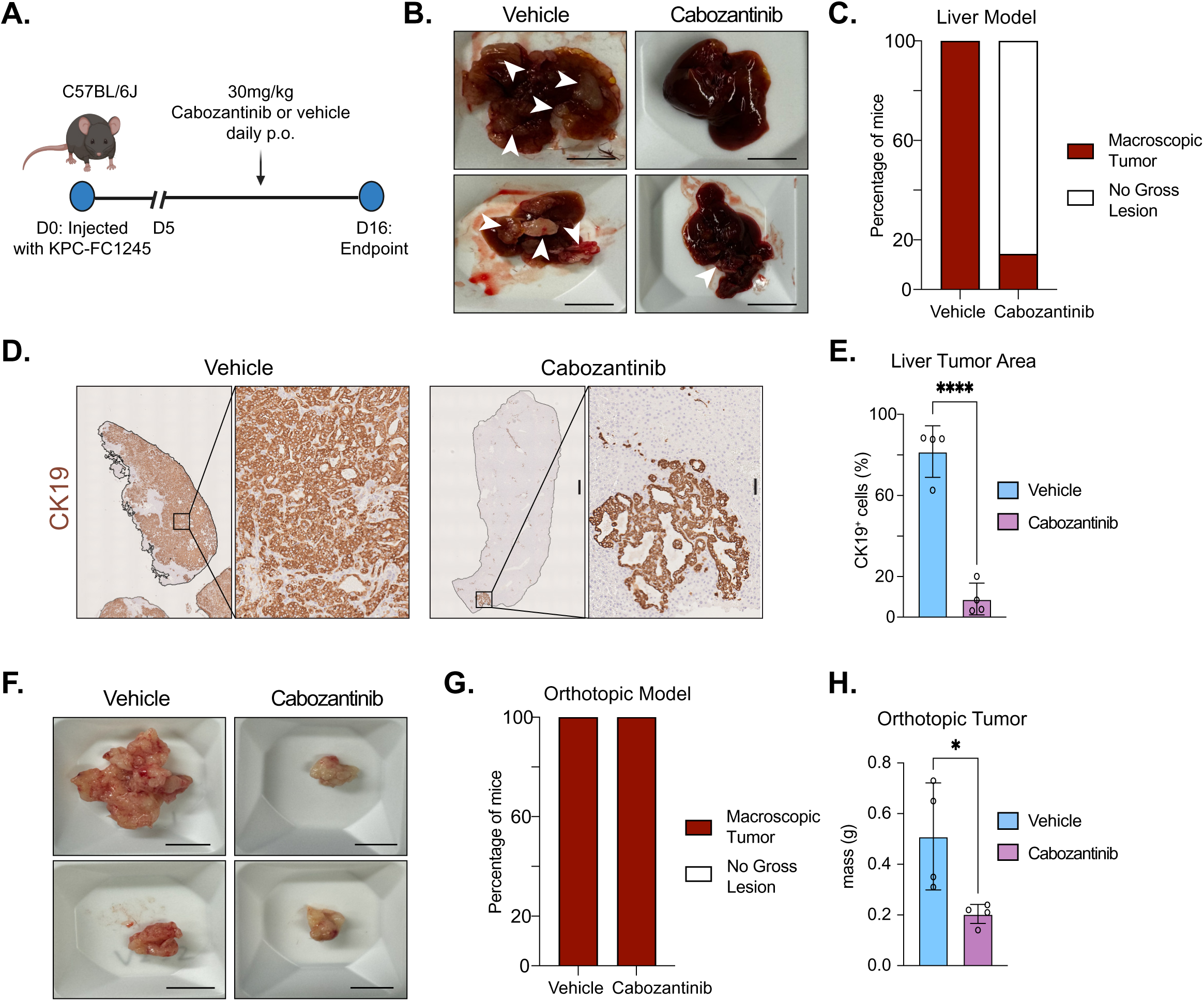
MET inhibition more potently impairs the growth of liver vs. pancreas PDAC. **A.** Schematic of experimental design for pharmacological MET inhibition. Here, KPC-FC1245 PDAC cells were implanted into the liver via hemi-spleen injections. 6 days later mice were randomly divided onto vehicle or 30mg/kg cabozantinib treatment arms p.o. daily for 12 days. **B.** Representative images of liver at endpoint from vehicle or cabozantinib treated mice. **C.** Quantification of the number of liver tissues that in (**B)** contained observable macroscopic tumors at harvest or appeared grossly normal. **D.** Representative IHC for Cytokeratin-19 (CK19) from liver tissue sections treated with vehicle or cabozantinib. **E.** Quantification of percent CK19^+^ cells per section, n= 4 per group. **F.** Representative images of pancreas tumors at endpoint from vehicle or cabozantinib treated mice. **G.** Quantification of the number of liver tissues that in (**F)** contained observable macroscopic tumors at harvest or appeared grossly normal. **H.** Mass of orthotopic tumors harvested from vehicle or cabozantinib treated mice. Scale bars are 1cM for gross tissues, 500µM for low magnification IHC, 50µM for high IHC magnification. Error bars are mean ±SD; * P ≤ 0.05; **** P ≤ 0.0001 by two-tail student’s t test.

Most pharmacological MET inhibitors, including cabozantinib also inhibit other MET family receptor tyrosine kinases including VEGFR and RET. To demonstrate the specific impact of targeting MET on PDAC liver tumor growth, we targeted MET expression in KPC FC1245 cells with CRISPR editing. Here, we were able to obtain a robust loss of MET expression in sg*Met* clonal cell lines (**Supplemental Fig. 7a**). We selected 4 sg*Met* clonal lines that individually grew slightly slower than the parental FC1245 control (**Supplemental Fig. 7b**), again demonstrating an important role for MET on PDAC biology. However, this defect was less obvious in a cocktail of the 4 sg*Met* clonal lines that we combined to use for in vivo studies.

We then established syngeneic liver tumors in mice C57BL/6J mice through hemispleen injections of sg*Met* KPC-FC1245 or parental controls (**Fig. 6a**). These mice were sacrificed and liver tissue harvested at Day 16. Here, we observed that the majority of sg*Met* tumors lacked gross macroscopic lesions that were observed in all the parental KPC-FC1245 tumors (**Fig. 6b,c, Supplemental Fig. 7c**). Assessing CK19 immunostaining, we observed that parental controls showed the liver tissue is largely replaced by PDAC cells, whereas the CK19^+^ sg*Met* tumors were either small or not seen (**Fig. 6d,e**).

**Figure 6:**
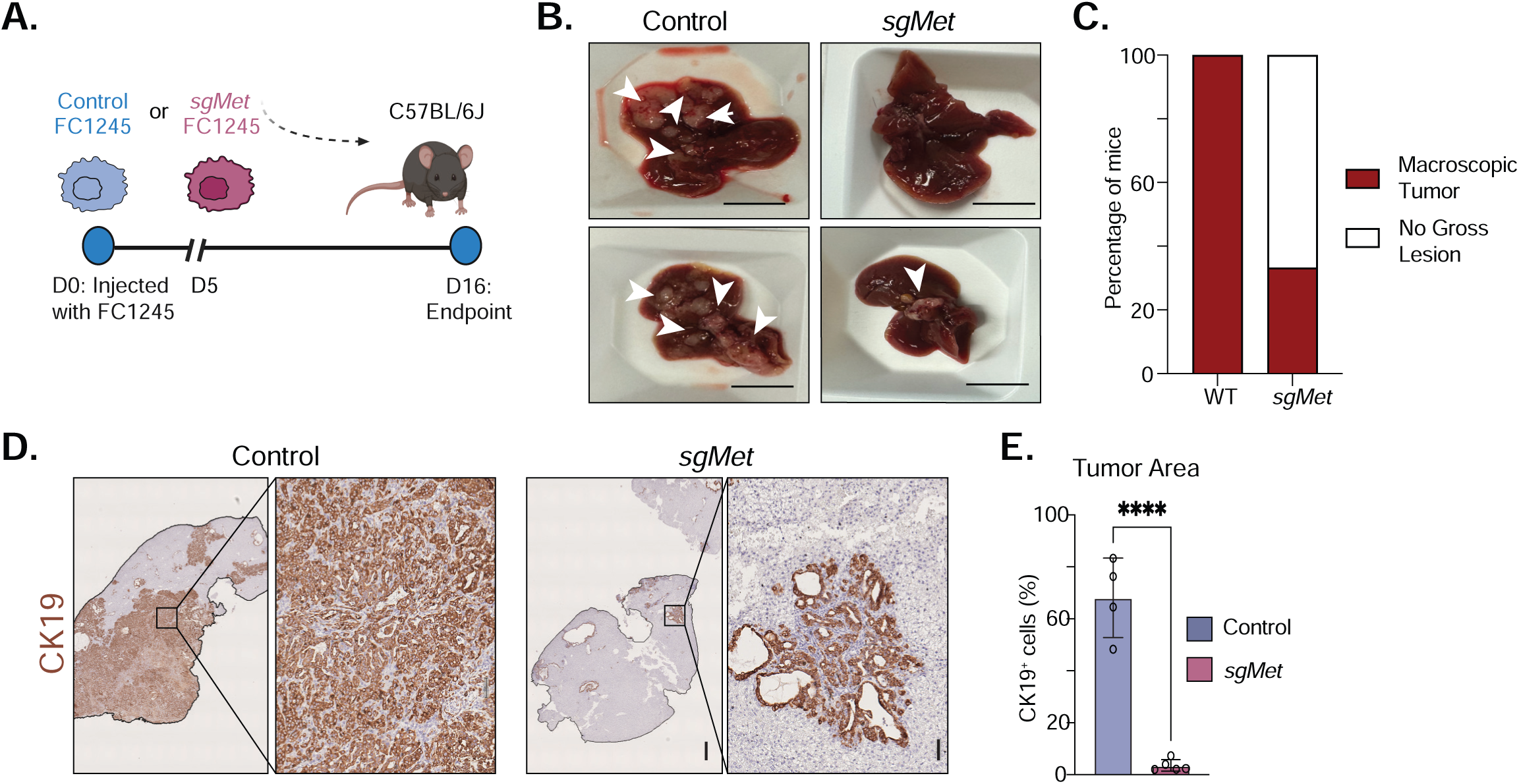
Genetic MET targeting recapitulates pharmacological inhibition. **A.** Schematic of experimental design for genetic *Met* targeting. Here, control or sg*Met* KPC-FC1245 PDAC cells were implanted into the liver via hemi-spleen injections and mice were sacrificed 16 days later. **B.** Representative images at endpoint of livers implanted with control or sg*Met* KPC cells. **C.** Quantification of the number of liver tissues that contained observable macroscopic tumors at harvest or appeared grossly normal. **D.** Representative CK-19 IHC staining of control or sg*Met* FC1245 implanted liver tissues. **E.** Quantification percent CK19^+^ cells per section (n=4 control, n=5 sg*Met*). Scale bars are 1cM for gross tissues, 500µM for low magnification IHC, 50µM for high IHC magnification. Error bars are mean ±SD; ** P ≤ 0.01; *** P ≤ 0.001; **** P ≤ 0.0001 by two-tail student’s t test.

Collectively, these data confirm that MET signaling is a key pathway promoting the growth of liver PDAC tumors. MET blockade of primary and liver PDAC tumors after engraftment results in an impairment of growth in both contexts. Importantly, this impact is more pronounced in the liver where both genetic and pharmacological inhibition results in most mice having no macroscopic tumors. Importantly, the MET pathway can be leveraged therapeutically using clinically available pharmacological inhibitors such as cabozantinib.

## Discussion

Desmoplasia has long been recognized as a characteristic feature of PDAC (39). Accordingly, pancreatic fibroblasts have been extensively studied to understand how they remodel the pancreas to support carcinogenesis and PDAC progression. These insights have led to better understanding of how CAFs coordinate with malignant epithelial and immune cells to create an environment permissive to PDAC growth (2). Among the first characterized stromal-PDAC crosstalk pathways was the activation of hedgehog signaling in CAFs by ligands released from malignant epithelial cells that promote stromal expansion and fibrosis (4,40). While disruption of this signaling axis has thus far proven disappointing in the clinical setting potentially either due to loss of a tumor restraining populations (6–8), or enrichment of more immune suppressive fibroblasts (41), this illustrates the central impact that CAFs mediate in PDAC biology.

The stromal populations in PDAC liver tumors have also been suggested to have tumor-promoting and restraining functions (29). In this study, we sought to directly compare how CAFs that arise in response to the same PDAC cells program liver lesions vs. pancreatic tumors. Here, we verified that the expansion of stroma associated with tumorigenesis in the pancreas is also observed in regions of the tumor-bearing liver. Using single cell RNA sequencing we observed that the vast majority of liver CAFs show transcriptional profiles distinct from those that map onto fibroblasts that occur in primary pancreatic tumors. Thus, although pancreatic fibroblasts have often been compared to liver fibroblasts, as stellate populations have been isolated from both organs, liver fibroblasts are clearly different from those found in the pancreas even when exposed to signals from the same cancer cells.

Importantly, this also indicates that stromal-cancer support networks between PDAC cells and CAFs are highly variable between tumor sites. Here, we observe that HGF expression is a feature of normal and cancer-associated fibroblast populations in the liver. Accordingly, the mitogenic signaling provided by MET activation in PDAC cells likely plays important roles in both the establishment and maintenance of the liver metastatic tumor niche. The role of HGF-MET signaling axis has emerged as a potential target in PDAC under the context of pancreatic stellate cells in primary tumors (37,42,43). Here, targeting HGF or MET was shown to potentiate response to several treatment approaches and inhibit metastasis of primary tumors. However, the focus on isolated pancreatic stellate cells in this study might overstate the impact of this axis in primary tumors as stellate-derived CAFs are a minority population in PDAC (9). HGF-MET signaling has also been identified as a signaling axis in liver iCAF populations, although the therapeutic potential remained unknown (29). Here, we confirm that the *HGF* expression in fibroblast populations from both human and mouse PDAC liver tumors and fibroblasts isolated from nearby normal human liver tissue is markedly higher than that found in primary pancreatic tumors. Accordingly, this HGF-MET crosstalk likely plays an important role in both establishment and maintenance of the liver metastatic niche.

As we sought to compare the interactions across many different populations of cells in pancreatic and liver PDAC lesions, the number of fibroblasts we captured from liver tumors occluded any confident in-depth characterization of potential subpopulations that likely co-exist within the CAFs present in the metastatic niche. However, recent studies probing the programming of CAFs in liver metastasis have suggested that *Hgf* is expressed by iCAFs or vascular-associated metastasis-associated fibroblasts (vMAFs) (29,30). Our data also supports the extensive literature of normal liver fibroblast HGF production in the context of wound-healing and tissue repair (44). We find liver-derived fibroblasts are able to activate canonical mitogenic signaling pathways downstream of MET including ERK, AKT, and STAT3 phosphorylation in PDAC cells in a MET-dependent fashion. Interestingly, we find that STAT3 is not activated in PATC53 PDAC cells by HGF alone and pharmacological inhibition of MET *in vitro* does not completely reduce ERK or STAT3 activity suggesting a potentiation of these pathway with other signaling molecules or metabolites released from liver fibroblasts (2). Despite this complexity, MET inhibition is clearly sufficient to potently impair the ability of PDAC cells to grow within the liver.

Single cell transcriptomics studies have nearly universally observed that PDAC sub-population heterogeneity within tumors does not correspond to the transcriptional subtypes defined from bulk RNA sequences approaches (2,45). In our datasets, we were able to define four main PDAC cell sub-populations with functionally relevant gene signatures that are present in both tumor sites. Interestingly, despite establishing tumors with the same initial population of PDAC cells, we observed a profound preference for programming centered on redox metabolism in the cells residing in the liver. This could be influenced by the stark differences in the metabolic state encountered between the hypoxic nutrient poor pancreatic tumor microenvironment (22,24) as compared to the liver, which serves as a primary anabolic hub and the center of gluconeogenesis. Indeed, metastatic PDAC cells have been shown to exhibit preferential metabolic pathway utilization(46), and this can serve to allow preferential seeding in one organ over another (47). Further, we have already identified that differential metabolic programming can co-exist in both human and murine PDAC (34), and recent work has shown clonal metabolic states are key to metabolic adaptation (35). In addition, we cannot conclude if there is plasticity between these populations or this over representation of redox metabolism preference in liver tumors is selected during seeding, although the dominance of specific clonal populations in PDAC metastasis has been reported using elegant genetic barcoding (48).

Finally, the sensitivity of PDAC cells to MET inhibition has been reported in the context of primary pancreatic tumors (42,49). Here, we used clinically available MET inhibitor cabozantinib to demonstrate the therapeutic potential in liver metastatic pancreatic cancer. This drug is currently approved to treat patients with renal cell carcinoma, thyroid cancer, and hepatocellular carcinoma where it has demonstrated clinical efficacy (50–52). Cabozantinib was also recently approved for the treatment of pancreatic neuroendocrine tumors after phase 3 trial showed 13.8 month of progression free survival compared to 4.4 months in the placebo group (53). Cabozantinib was explored in combination with gemcitabine in a small cohort of PDAC patients and showed some response that would warrant further investigation (54). However, dose limiting toxicities were an issue in this phase 1 trial and a maximum tolerated dose could not be obtained. For example, combining cabozantinib with immunotherapy or other targeted agents may offer a less toxic therapeutic strategy. This approach is currently under evaluation in clinical trials in PDAC, including cabozantinib + atezolizumab (NCT04820179), cabozantinib + pembrolizumab (NCT05052723), and cabozantinib + erlotinib (NCT03213626). Accordingly, while there is potential for MET inhibitors to be used in PDAC, the right contexts will need to be found and tested in further trials. Given our observations, we expect that cabozantinib could be an ideal application as a first line single agent or in combination regimens with other targeted agents as chemotherapy is unlikely to have a durable response in PDAC liver metastasis. Further, given the frequency of recurrence in the liver, after curative intended tumor resection, a sequential treatment strategy such as adjuvant treatment cabozantinib following standard of care chemotherapy may be effective to eliminate disseminated cells before they can establish a liver tumor niche.

## Acknowledgements

We would like to thank Devon Pendlebury and the members of the Halbrook Lab for support and critical reading of the manuscript, Delia Tifrea and the UCI Experimental Tissue Repository for specimen acquisition, Dr. Maksim Plikus for advice on CellChat analysis, Dr. Marcus Seldin, Himanshu Gupta, and Gregory Tong for computational support. C.J.H. was supported by R00CA241357, R37CA283575, American Cancer Society Institutional Research and Research Scholar Grants (RSG-1255258), a V Scholar award (V2021-026), a UCI Anti-Cancer Challenge Pilot award, and a Tower Cancer Research Foundation Career Development Award. C.J.H, T.F.M, D.I, and Z.J, were supported by the Chao Family Comprehensive Cancer Center support grant (P30CA062203). A.K.B was supported by a fellowship through the German Cancer Aid Foundation (Mildred-Scheel-Postdoktorandenstipendium). O.G.M was supported by R01 CA222594. T.F.M. was supported by K01CA249038. N.G.S. was supported by R00CA263154.

## Author Contributions

CJH and RS conceived of, designed this study, and wrote the manuscript. CJH, NY, CA, AKB, WMA, SC, IAL, OGM, JPM, DI, ZJ, TFM, NGS provided key reagents, performed experiments, analyzed, and interpreted data. CJH supervised the work carried out in this study and obtained funding.

## Methods

### Study approvals

All procedures were performed at the University of California, Irvine in compliance with the Institution Animal Care and Use Committee (AUP-23-084) and the Institutional Biosafety Committees (BUA-R315). Patient specimens were obtained de-identified through UCI 08-70 or the UCI Experimental Tissue Repository.

### Cell culture

The KPC-FC1245 cell line was a gift from Dr. David Tuveson (Cold Spring Harbor Laboratory). PATC53 cells were obtained from ATCC. PDAC cells were maintained in high-glucose DMEM (Gibco) supplemented with 10% FBS (Corning). Primary fibroblasts were established by outgrowth from tissue pieces as described (16) and maintained in DMEM with 20% FBS. All cells were routinely tested for mycoplasma contamination using MycoAlert PLUS (Lonza). Cabozantinib was obtained from ChemGood. HGF was obtained from Peprotech.

### Orthotopic surgery

Orthotopic transplantation into the pancreas was performed using 50,000 KPC FC1245 cells prepared in a 1:1 ratio with Matrigel and media (DMEM + 10% FBS). Mice were anesthetized with isoflurane and area for surgery was prepared using aseptic techniques. A 2-inch incision was made subcutaneously and intraperitoneally, then 50uL of tumor cell suspension was injected into the tail of the pancreas.

### Hemi-spleen surgery

For liver tumor seeding, the spleen was accessed through a Laparotomy then expressed through the skin. The spleen was then ligated into two halves inferior to the hilar vessels using medium sized ligating clips and a clip applicator. After making an incision on the spleen, the lower half was tucked back into the peritoneum. A cell suspension with KPC FC1245 cells followed by a layer of PBS were loaded in a syringe and injected at a 30-degree angle right underneath the spleen capsule. After injecting the full volume, the needle was left inside the spleen to avoid any fluid escape. Another ligation clip was placed to ligate the superior hilar vessels and the remaining spleen was removed using scissors.

### Murine Single Cell Transcriptomics

Tumors were removed and minced using a scalpel. Further dissociation was achieved with enzymatic digestion with collagenase V for pancreas tumor and collagenase IV for liver tumors (1mg/mL DMEM) for 30 mins on a MACS tissue dissociator. Tumor suspension was then filtered through 500-um, 100-um, and 40-um mesh to obtain single cells. Dead cells were subsequently removed using MACS Dead Cell Removal Kit (Miltenyi Biotec.). Single cell DNA libraries were prepared and sequenced at the UCI Genomics Research and Technology Hub. Cell Ranger version (chemistry 3’ v3, pipeline version 3.1.0) was used with default settings with an initial expectation of 10,000. Cell Ranger software was used for alignment and quantification. For downstream analyses, R studio version 4.1.1 was used. Seurat version (5.2.1) was used as previously described (55). Briefly, QC was performed to exclude cells containing less than 200 or more than 2000 genes. Data were normalized using LogNormalize with a scale factor of 10,000 followed by scaling. PCA was run on the filtered data. Clusters were defined based on the gene expression using FindAllMarkers. The code is publicly available on GitHub (https://github.com/halbrook/HalbrookLab).

### Human Single Cell Transcriptomics

Human single cell RNA sequencing atlas data was utilized from the following prior study (Loveless et al., Clin Can Res, 2024). Briefly, this atlas included 172 primary tumors and 25 metastatic biopsies, the majority of which were from liver metastatic PDAC tissues. Tissues were a mix of treated and untreated samples. Raw data were aligned to the same human reference genome and batch correction was performed with Harmony package. Seurat v4 was used to cluster data and cell types were assigned based on major lineage markers such as *KRT18/19* for ductal identify, and *DCN, LUM, COL1A1* for CAFs. Ductal cells and CAFs were clustered separately, and DotPlot visualizations were used for specific genes of interest. Classical, basal, or hybrid identify in ductal subpopulations was determined by scoring of gene signatures from a prior study (Raghavan et al., Cell).

### Histology

Mice were sacrificed by CO2 asphyxiation then tissue was quickly harvested and fixed overnight at room temperature with Z-fix solution (Anatech LTD). Tissues were processed using a Leica ASP300S Tissue Processor, paraffin embedded, and cut into 5 µm sections. Immunohistochemistry was performed by deparaffinizing and rehydrating tissues then performing antigen retrieval with a Sodium Citrate Buffer (10mM Sodium Citrate, 0.05% Tween 20, pH 6.0). Blocking was performed with serum free protein block (Dako), then PDGFRβ (Abcam, #32570, 1:300) and Cytokeratin 19 antibody (Abcam, #ab133496, 1:500) were used as primary antibodies. After washing, Rabbit-on-Rodent HRP Polymer (Biocare Medical) was used as secondary antibody, then chromogen deposited using DAB plus (Dako). Slides were then counterstained with hematoxylin. Hematoxylin and eosin staining were performed per manufacturer’s instructions. Immunohistochemistry for PDGFRβ (Abcam, #32570) or MASP1 (ThermoFisher, #PA5-47992) was performed separately with standard conditions on a Discovery ULTRA stainer (Roche) and antibodies were detected with DAB, and counter stained for nuclear detection. Brightfield images were taken on a Leica THUNDER microscope.

### Fluorescence ISH

Five-micron sections were cut from adjacent or pancreatic cancer metastatic liver biopsy tissues onto charged slides. Slides were then baked at 60°C for 30 minutes, deparaffinized with xylene for 10 minutes, dehydrated in 100% ethanol for 2 minutes, and washed with 0.1% Tween-20 RNAse-free 1x phosphate-buffered saline (PBST). RNA scope Multiplex Fluorescent Detection v2 kit assay was performed according to manufacture instructions (Advance Cell Diagnostics; ACD). Briefly, slides were incubated with hydrogen peroxide for 10 minutes at room temperature followed by target retrieval at 98°C for 15 minutes. Slides were then blocked with the Co-Detection antibody diluent for 30-60 minutes and incubated with Pan cytokeratin (PanCK) (Invitrogen, #53-9009-82, 1:400) for 16 hours at 4°C. The following day tissue sections were post-fixed with neutral buffered formalin and treated with the ProteasePlus Reagent for 13 min at 40°C in a hybridization oven. Amplification and signal enhancement (AMP) were performed for either a single probe (C1) or two different probe (C1 and C2) combinations. UPP1-C1 was a 1x probe and DCN-C2 was a 50x probes. Probes were diluted in ACD probe diluent per manufacturer’s instructions and slides were incubated with them at 40°C for 2 hours. Following two washes with RNA scope washing buffer, the signal for each probe was amplified with AMP reagents, horseradish peroxidase, and tyramide signal amplification kit at 40°C. Slides were then incubated with anti-mouse or anti-rabbit secondary Alexa Fluor 488 IgG (H + L) antibody (1:400) for 1 hour at room temperature. Tissue sections were counterstained with DAPI for 15 minutes at room temperature and washed three times with PBST before being mounted with ProLong Diamond Antifade mounting medium. Images were taken on a white light laser confocal STELLARIS (Leica) microscope.

### Immunofluorescence

Immunofluorescence for MASP1 (ThermoFisher, #PA5-47992, 1:100), PDGFRβ (Abcam, #32570, 1:300), or αSMA (Sigma, #A2547, 1:1000) was performed as previously described (Steele et al., Nature Cancer, 2020). Briefly, slides were deparaffinized and underwent citric acid retrieval in a microwave for 20 minutes. After cooling slides were blocked in 20% donkey serum for 30 minutes at room temperature. Following this, primary antibodies were added at 4°C for 16 hours. Tissue sections were counterstained with DAPI for 15 minutes at room temperature and washed three times with PBST before secondaries were added (Jackson Immuno donkey secondaries at 1:400 in PBS were used against goat, mouse or rabbit) for 45 minutes at room temperature and then slides were mounted with ProLong Diamond Antifade mounting medium. Images were taken at 20x or 40x on a white light laser confocal STELLARIS (Leica) microscope. Images were quantified with HALO software and at least 5 large-scale regions (N=3 Primary PDAC, N=4 Liver metastasis) were analyzed for each 40X tile scanned image (average number of cells per high power field were averaged for each patient within these regions).

### CellChat Network Analysis

Differential cell–cell communication analysis was conducted using the CellChat framework. Specifically, intercellular communication probabilities between each cell-type pair were inferred independently for liver and pancreas tumor, utilizing the curated interaction database provided by CellChatDB. Subsequently, differential communication strength between the two conditions was computed for each cell-type pair to identify context-specific alterations in signaling networks.

### Bulk RNA Sequencing

Primary fibroblasts and CAFs were brought up in culture and upon reaching 70-80% confluency, RNA was harvested using RNeasy Plus Mini kit (Qiagen, #74126). RNA quality was verified using Nanodrop. Samples were sent for sequencing. Quality was ensured using RIN (above 8). FastQFiles were then aligned using STAR, bedgraphs were generated, and RNA counts were obtained in tpm format. Downstream analyses were performed in R studio. Briefly, differential expression of genes was performed to generate Venn diagram and z-scores were obtained to plot heatmaps.

### ELISA

Serum free conditioned media was harvested from primary fibroblasts. ELISA was performed using the Human HGF Quantikine ELISA kit (R&D, #DHG00B) and following manufacturer’s instructions.

### Western Blotting

Cells were lysed with 1X RIPA lysis buffer (Sigma-Aldrich Cat# 20188) supplemented with protease inhibitors (cOmplete™, EDTA-free Protease Inhibitor Cocktail, Sigma-Aldrich Cat# 11873580001) as well as phosphatase inhibitors (PhosSTOP™, Roche Cat# 4906845001). Protein quantification of cleared lysates was performed using a bicinchoninic acid (BCA) assay kit (Pierce™ BCA Protein Assay Kits, ThermoFisher Cat# 23227). Lysates were subsequently electrophoresed and immunoblotted with the indicated primary antibodies: Phospho-Met (Tyr1234/1235) (1:1000, Cell Signaling Technology Cat# 3077T); Met (1:1000, Cell Signaling Technology Cat# 3127); p44/42 MAPK (Erk1/2) (1:1000, Cell Signaling Technology Cat# 4695S); Phospho-p44/42 MAPK (Erk1/2) (Thr202/Tyr204) (1:1000, Cell Signaling Technology Cat# 4370S); β-Actin (1:5000, Santa Cruz Biotechnology Cat# sc-47778); Vinculin (1:5000, Cell Signaling Technology Cat# 13901S). Nitrocellulose membranes(Bio-Rad) were then incubated with the appropriate HRP-conjugated secondary antibodies: Anti-rabbit IgG, HRP-linked Antibody (1:10,000, Cell Signaling Technologies Cat# 7074S); Anti-mouse IgG, HRP-linked Antibody (1:10,000, Cell Signaling Technologies Cat# 7076S); or IRDye 800CW Goat anti-Mouse IgG Secondary Antibody (1:15,000, Licor, 926-32210). Imaging was performed on a Thermo iBright Imaging System using an enhanced chemiluminescence (ECL) kit (SuperSignal™ West Femto Maximum Sensitivity Substrate, Thermo Fisher Cat# 34095) or on a Li-Cor Odyssey CLx.

### Cell Signaling Assays

PATC53 and PATC53 sgMet cells were grown in DMEM+10% FBS (Gibco) they reached 75% confluence. PATC53 plates were then switched to serum free DMEM for 12 hours prior to the assay. Serum free media was aspirated and plates were washed with PBS (Gibco) then subjected to treatments. Here, cells were treated with 1µm cabozantinib (ChemGood Cat# C-1333) or vehicle (0.5% DMSO) for 20 min then treated with 1ng/mL human HGF (Peprotech Cat#100-39-10UG) or vehicle (water) then incubated at 37°C for the times indicated in the figure legend for each assay.

#### Condition Media Assays

Liver fibroblasts, Liver CAFS, and PDAC CAFs were grown in DMEM+10% FBS (Gibco) they reached 75% confluence. Cells were washed with PBS then serum-free DMEM was added. Media was allowed to condition for 48 hours, then filtered through a 0.45µM filter. PATC53 and PATC53 sgMet cells were grown in DMEM+10% FBS (Gibco) until they reached 75% confluence. PATC53 plates were then switched to serum free DMEM for 12 hours prior to the assay. Serum free media was aspirated and plates were washed with PBS (Gibco) then incubated in serum free DMEM at 37°C. After 20 minutes, DMEM or 3:1 conditioned media – serum free DMEM was and incubated at 37°C for the time indicated in each figure legend. The conditioned media+Vehicle condition was incubated with DMEM+0.5% DMSO for 20 minutes at 37°C. For Cabozantinib treatment experiments, 1µm drug (or 0.5%DMSO) was added for 20 minutes at 37°C prior to treatment with 3:1 conditioned media or fresh DMEM.

#### CRISPR/Cas9 editing of PDAC Cell lines

Genetic targeting of MET in PDAC cell lines was achieved using CRISPR/Cas9 method described previously (23). Briefly, sgRNA sequences targeting *MET* or *Met* were selected from the human and mouse GeCKOv2 CRISPR knockout pooled library, respectively. Overlapping oligonucleotides were purchased, annealed, phosphorylated, then ligated into the overhangs of PX459 V2.0 vector (Addgene plasmid #62988) that was digested with BbsI. The resulting CRISPR/Cas9 plasmid was transformed into chemically competent Stbl3 cells, miniprepped for plasmid DNA, and sequence-verified. sgRNA oligonucleotide pairs hMET-PX459F CACCgCACATGGCAGATCGATCCAT and hMET-PX459R AAACATGGATCGATCTGCCATGTGc or mMET-PX459F CACCgCACATGGCAGATCGATCCAT mMET-PX459R AAACTGAATAAGTCGACGCGCTGCc were used. PDAC cells were transiently transfected using Lipofectamine 3000 according to the manufacturer’s instructions. Cells were selected with 2µg/mL puromycin for 72 hours, where a parallel non-transfected control plate was observed to be completely killed. To select clones, polyclonal pools were seeded into 96-well plates at a density of 1 cell per well. Individual clones were expanded and editing efficiency assessed via western blot.

### Mouse tumor treatments

C57BL/6J mice were injected with KPC FC1245 (250k/mouse). Tumors were established for 6 days before mice were randomized into vehicle or treatment arms. Mice (n= 5-7 per group) were treated with vehicle or cabozantinib (ChemGood #C-1333) (30mg/kg) every day for 12 days via oral gavage. After 12 days of treatment, mice were sacrificed and liver tissues were harvested upon sacrificing the mice.

### Data availability

scRNA sequencing data and bulk RNA sequencing data will be uploaded to GEO upon acceptance of manuscript. Human single cell RNAseq atlas is uploaded to Zenodo (https://zenodo.org/records/14199536). Other data will be provided upon reasonable request.

### Statistics & Reproducibility

All experiments were run a minimum of two times with at least 3 biological replicates. Statistics were performed using Graph Pad Prism 8 (Graph Pad Software Inc). Groups of 2 were analyzed with two-tailed students t test, groups greater than 2 were compared using one-way ANOVA analysis with Tukey post hoc test. All error bars, group numbers, and explanation of significant values are presented within the figure legends. The following values are used to denote significance; * P ≤ 0.05; ** P ≤ 0.01; *** P ≤ 0.001; **** P ≤ 0.0001. Sample sizes were determined by previous experiments performed in our groups. No data were excluded from the analyses.

