## Supplemental Figures for "Comparative Analysis of Primary and Liver Fibroblasts Reveals MET as a Potent Target in Pancreatic Cancer Metastasis"

### Supplementary Fig. 1: Characterization of cell populations in PDAC lesions

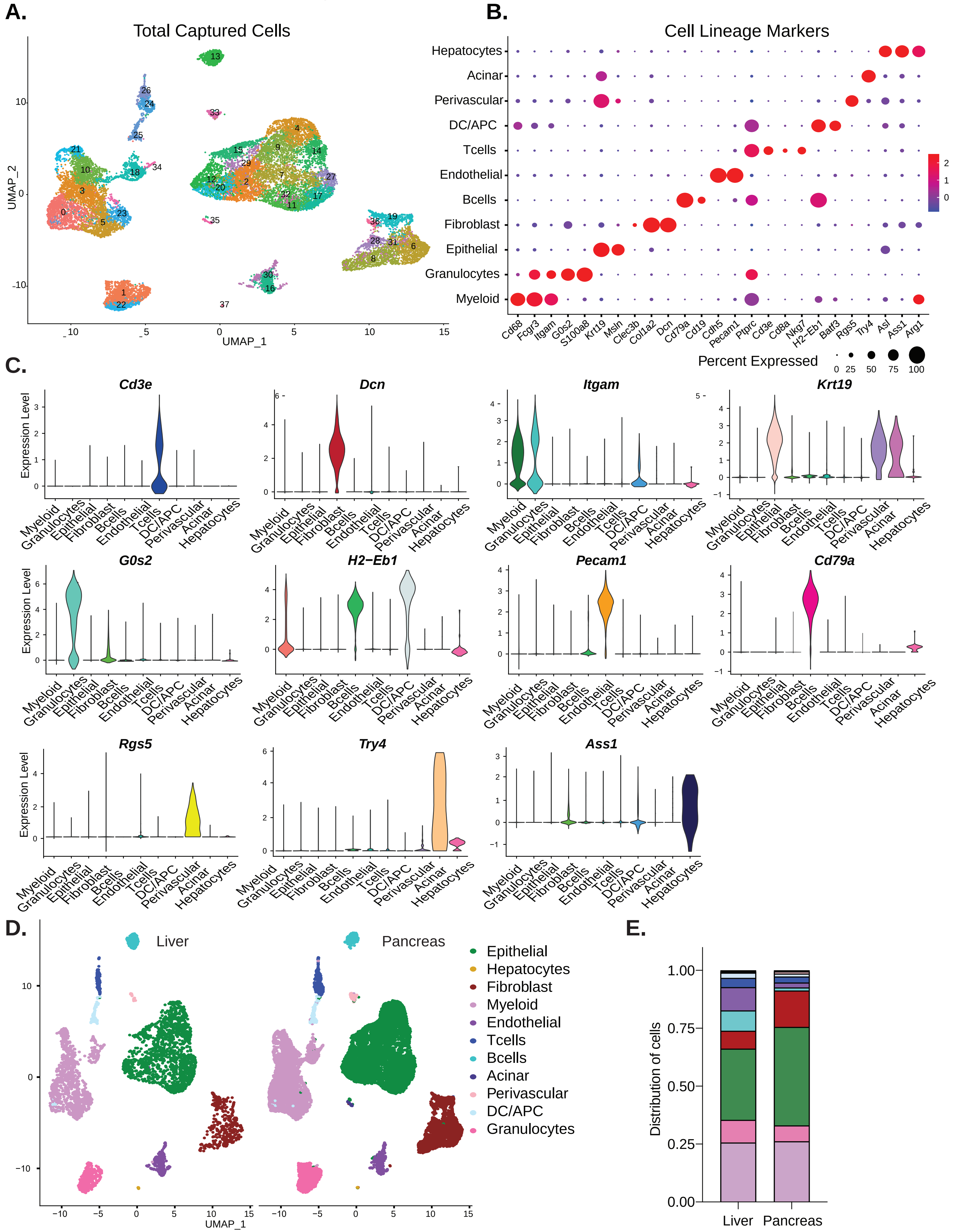

**Supplemental Figure 1:**

***Characterization of cell populations in PDAC lesions.*** **A.** UMAP plot of 38 different clusters calculated by Seurat representing all cells captured from single cell RNA sequencing of murine pancreas and liver PDAC tumors. **B.** Dot plot representation of marker genes used to define clusters in (**A**). **C.** Violin plots validating key defining genes- *Cd3e, Dcn, Itgam, Krt19, G0s2*, *H2-Eb1, Pecam1, Cd79a, Rgs5, Try4,* and *Ass1* used to identify each population. **D.** UMAP split by tumor site of identified cell lineages. **E.** Fractional distribution of cell populations between liver and pancreas tumors.

### Supplemental Fig. 2: MASP1 is expressed in liver, not pancreatic, fibroblasts

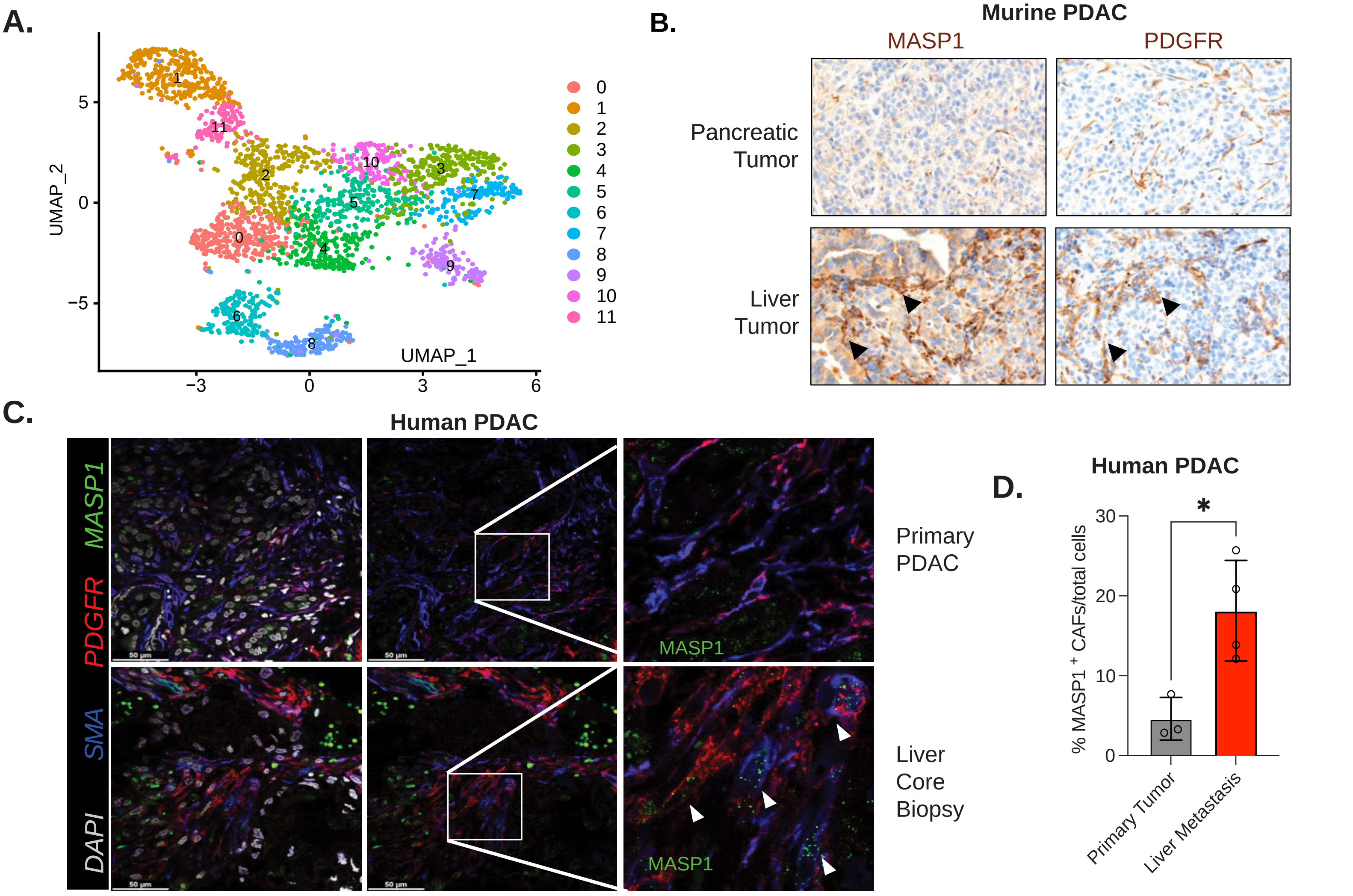

**Supplemental Figure 2:**

***MASP1 is expressed in liver, not pancreatic, fibroblasts.*** **A.** UMAP plot of 12 different clusters of PDAC fibroblasts. **B.** IHC staining on serial murine PDAC liver and pancreas tumors for the pan-fibroblast marker PDGFR and liver CAF marker MASP1. **C.** Representative co-immunofluorescence staining for MASP1 (green), PDGFR (red), fibroblast marker SMA (blue), and DAPI (white) in primary human PDAC tumor and metastatic liver core biopsies. **D.** Quantification of $MASP1^+$ fibroblasts in primary human PDAC vs. liver metastasis, n= 4 per group, ** P ≤ 0.05 by two-tailed student's t test.

### Supplemental Fig. 3: Murine PDAC cell clustering and human correlation

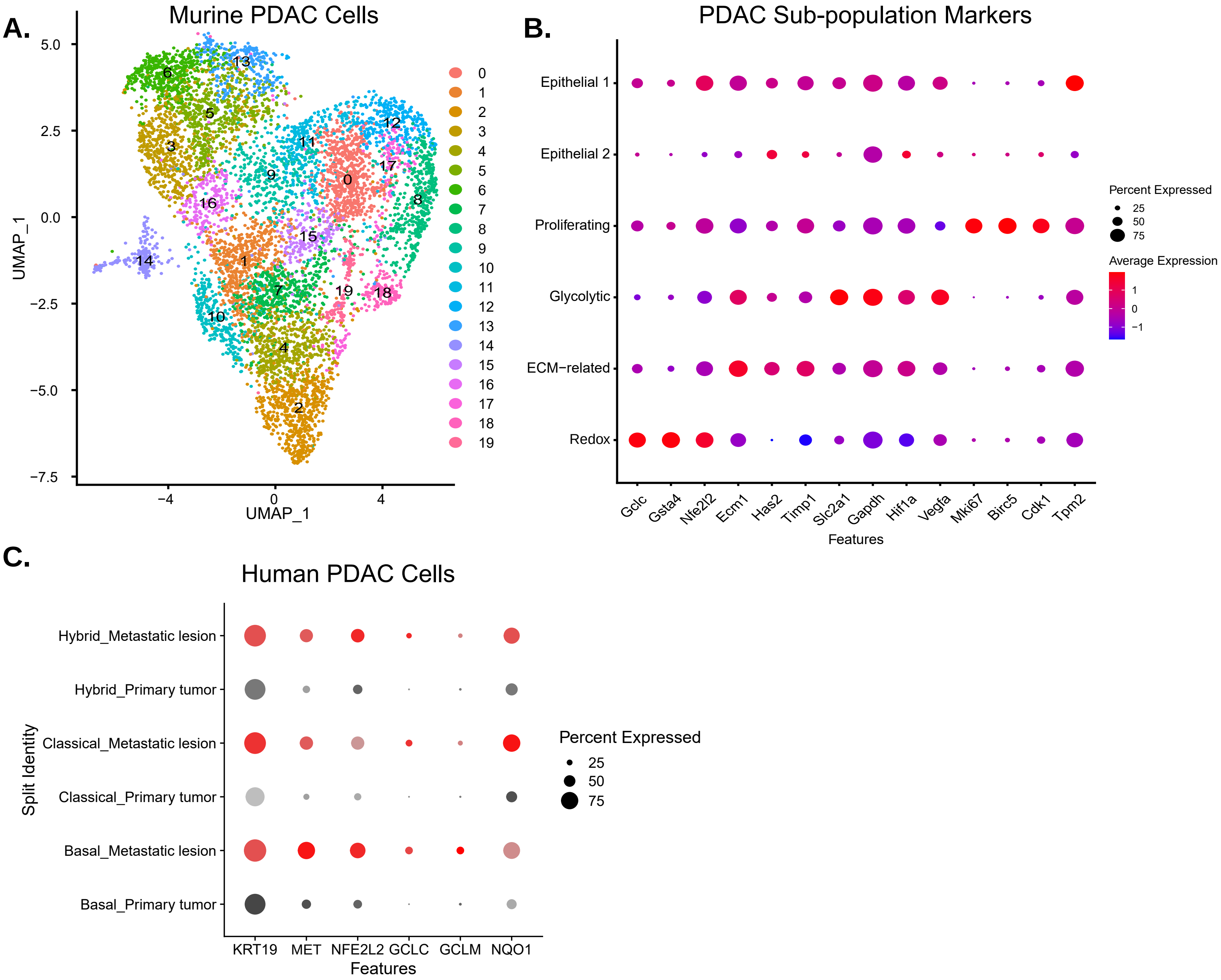

**Supplemental Figure 3:**

***Murine PDAC cell clustering and human correlation.*** **A.** UMAP of the 20 clusters of the PDAC cell object. **B.** Dot plot representation of key genes that were used to collapse clusters and denote sub-populations. **C.** Dot plot representation of redox metabolism genes *NFE2L2, GCLC, GCLM*, and *NQO1* in *KRT19* expressing human PDAC cells separated into Basal, Classical, or Hybrid transcriptional subtypes.

### Supplemental Fig. 4: HGF is not a feature of the pancreatic tumor signaling network

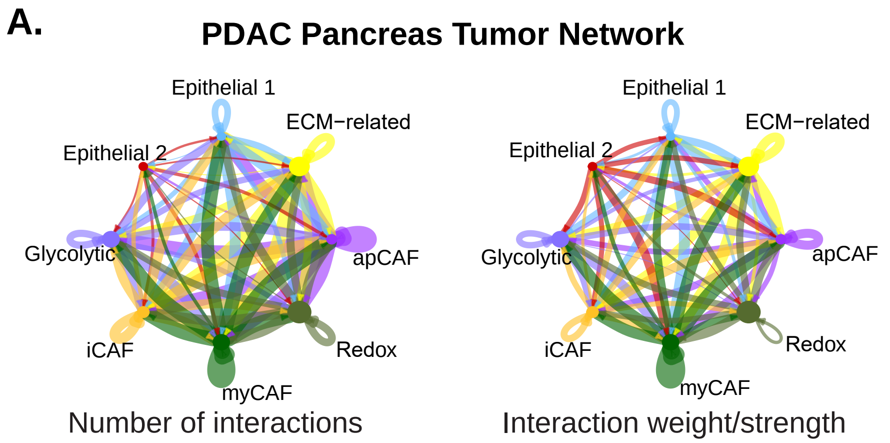


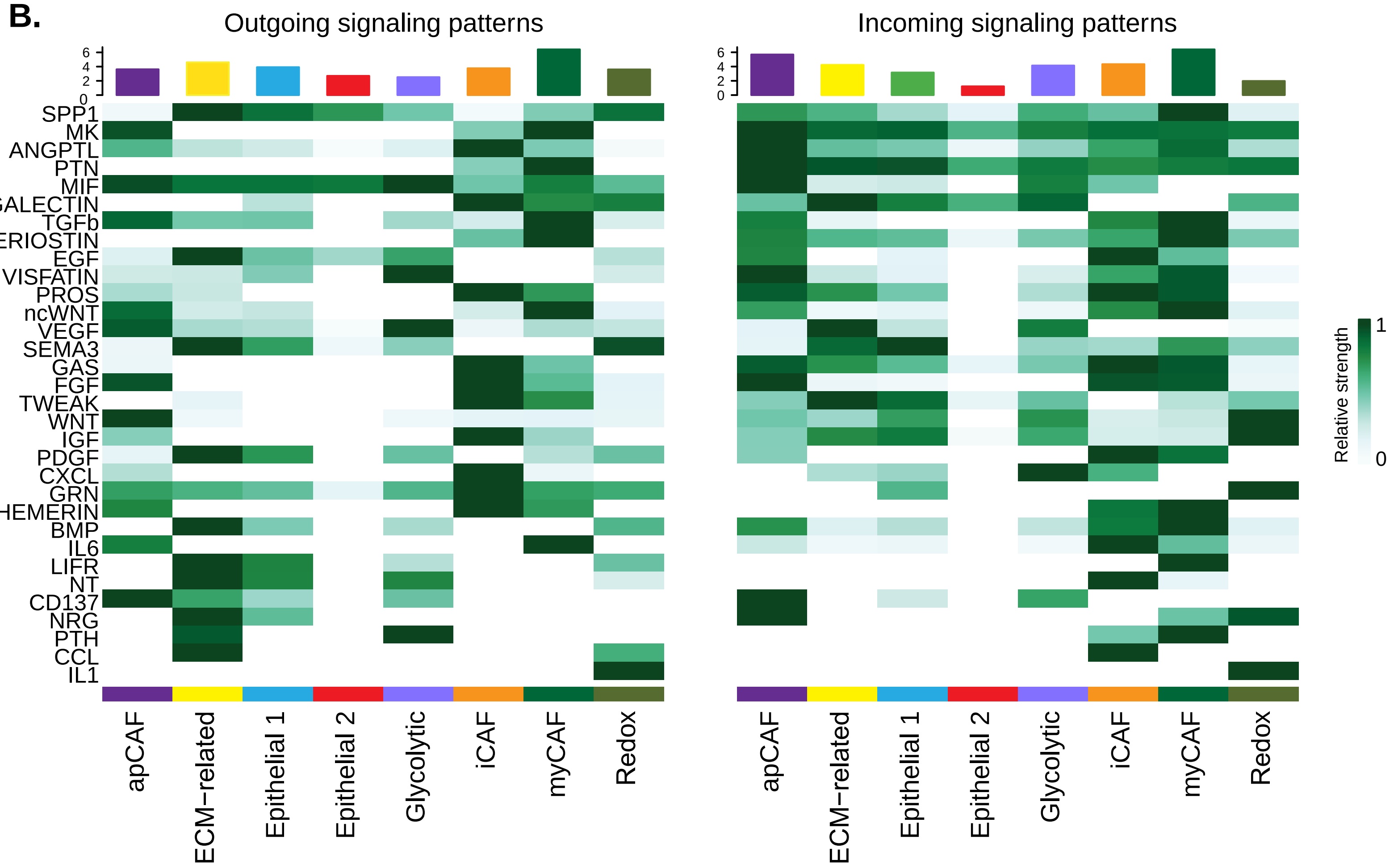

**Supplemental Figure 4:**

***HGF is not a feature of the pancreatic tumor signaling network***. **A.** Circle plot depiction of the CellChat aggregated cell-cell communication network showing number of ligand-receptor interactions (left) and strength of these interactions (right) between each indicated two populations in murine PDAC pancreatic tumors. **B.** Top signaling pathways with the most outgoing or incoming signaling in PDAC pancreatic tumors.

### Supplemental Fig. 5: Fibroblast Transcriptomics and sg*MET* PDAC signaling

A.

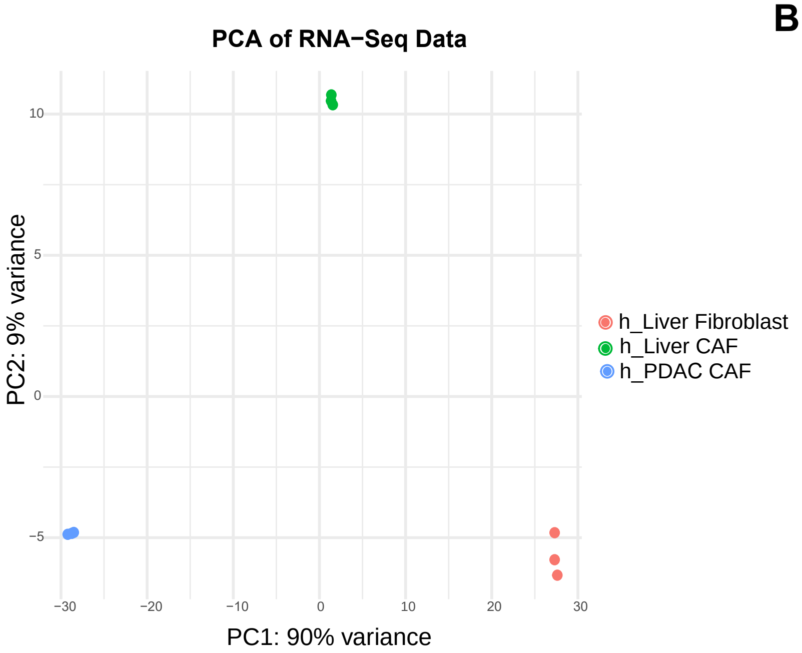


B.

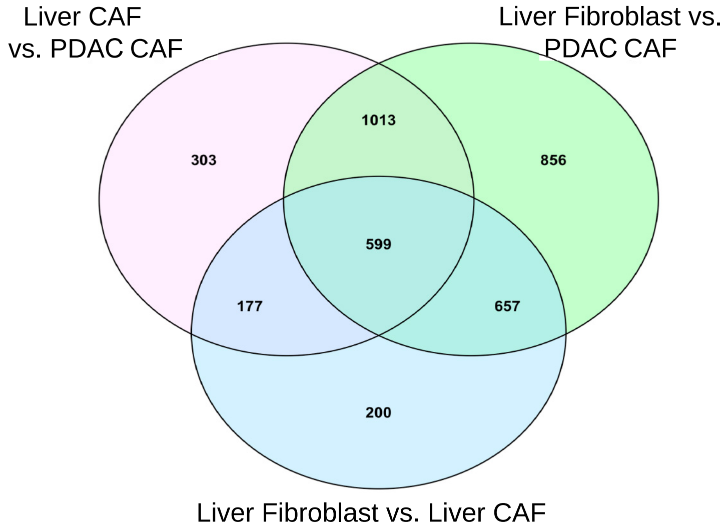


C.

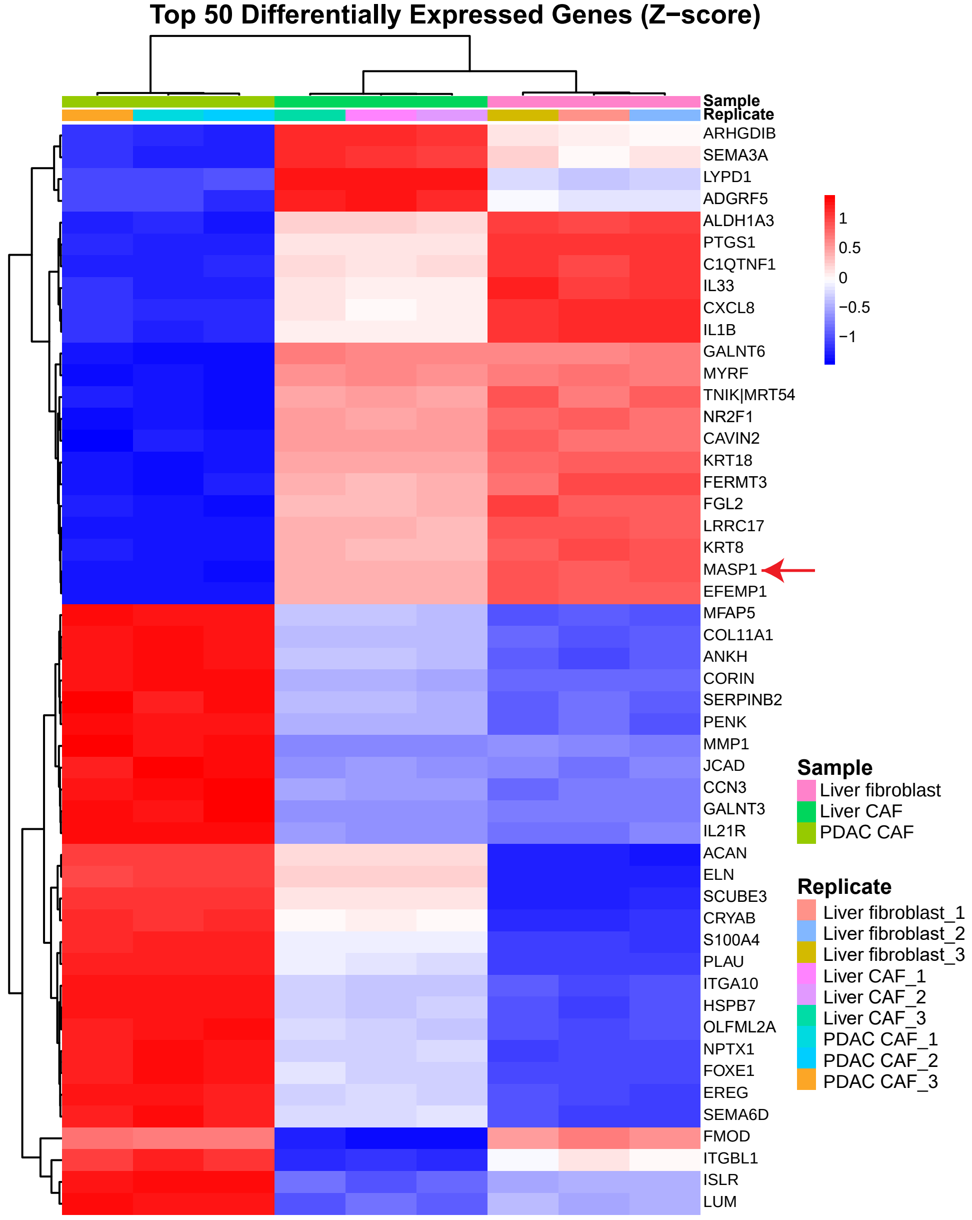

**Supplemental Figure 5:**

***Fibroblast Transcriptomics.*** **A.** Principal Component Analysis (PCA) performed on RNA sequencing data of human PDAC CAFs, Liver CAFs, and Liver Fibroblasts. **B.** Venn diagram depiction of the upregulated genes identified by differential expression gene analyses between the three fibroblasts lineages. **C.** Heat-map showing top 50 differentially expressed genes across all samples and groups, highlighting *MASP1* in RED.

### Supplemental Fig. 6: Cabozantinib data supplements

**A.**

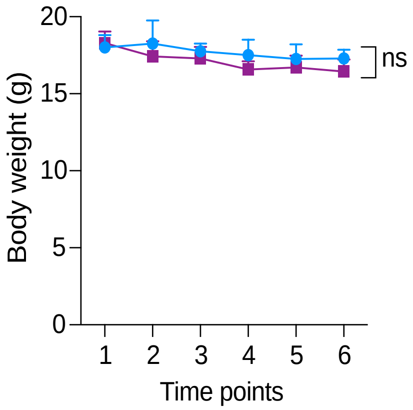


**B.**

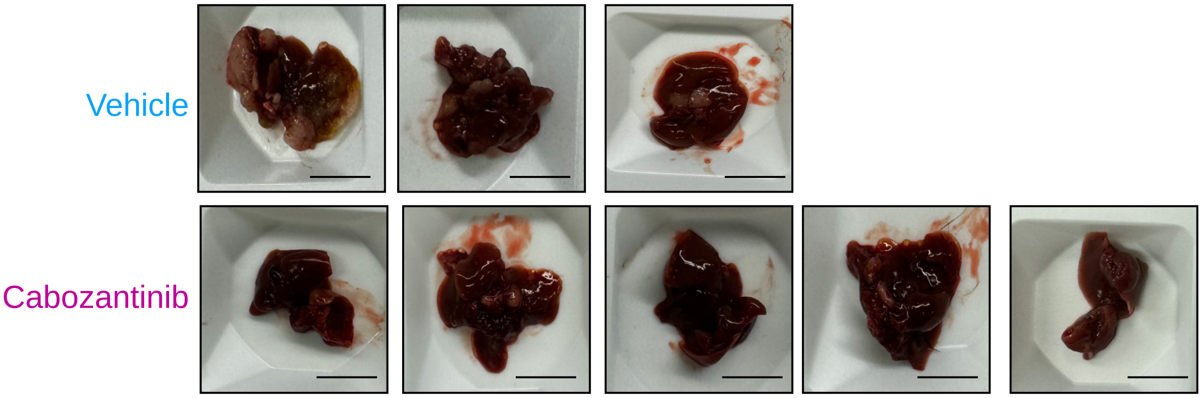


**C.**

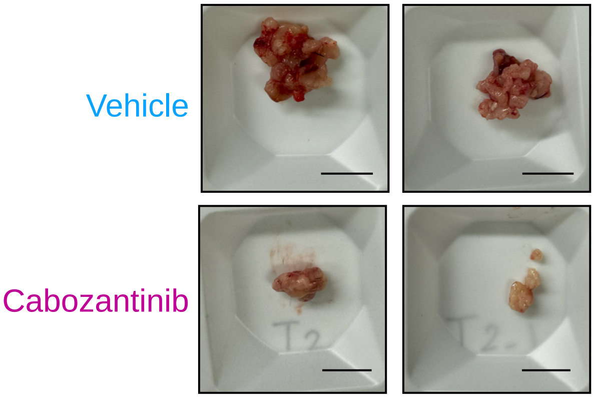

**Supplemental Figure 6:**

***Cabozantinib data supplements*. A.** Body weight of vehicle and cabozantinib treated mice throughout the experiment outline in **Figure 5a**. **B.** Gross liver images of remaining vehicle and cabozantinib treated mice harvested at the endpoint of the experiment. **C.** Gross pancreas images of remaining vehicle and cabozantinib treated mice harvested at the endpoint of the experiment.

### Supplemental Fig. 7: *sgMET* validation and data supplements

A.

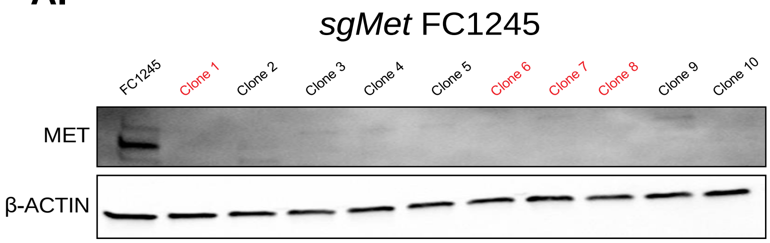


B.

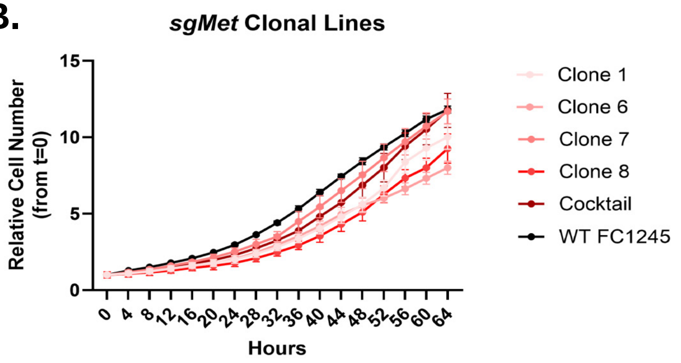


C.

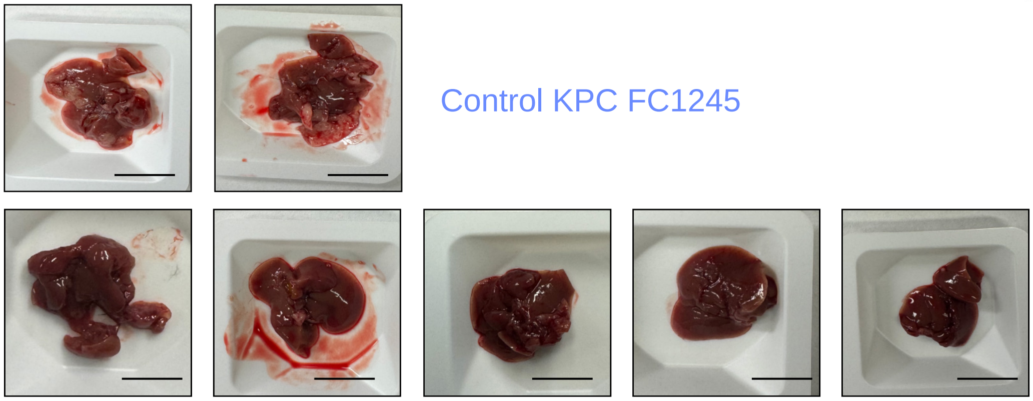

Control KPC FC1245

sgMET KPC FC1245

**Supplemental Figure 7:**

***sgMet validation and data supplements.*** **A.** Western blot showing extent of MET loss in 10 different clonal sg*Met* KPC FC1245 lines vs. parental control, later pooled clones highlighted in red. β-ACTIN was used as the loading control. **B.** Growth curves of 4 selected sg*Met* FC1245 clones, the 4 clones pooled together (cocktail), and parental FC1245 cell line as relative cell number (from t=o) over 64 hours analyzed by Cytation5 imaging. **C.** Gross liver images of mice injected with control FC1245 or sg*MET* FC1245 taken at the endpoint.
